# Gut microbe *Lactiplantiballus plantarum* undergoes different evolutionary trajectories between insects and mammals

**DOI:** 10.1101/2022.03.04.482961

**Authors:** Elisa Maritan, Marialaura Gallo, Dagmar Srutkova, Anna Jelinkova, Oldrich Benada, Olga Kofronova, Nuno F. Silva-Soares, Tomas Hudcovic, Isaac Gifford, Jeffrey E. Barrick, Martin Schwarzer, Maria Elena Martino

## Abstract

Animals form complex symbiotic associations with their gut microbes, whose evolution is determined by an intricate network of host and environmental factors. In many insects, such as *Drosophila melanogaster*, the microbiome is flexible, environmentally determined and less diverse than in mammals. In contrast, mammals maintain complex multispecies consortia that are able to colonize and persist in the gastrointestinal tract. Understanding the evolutionary and ecological dynamics of gut microbes in different hosts is challenging. This requires disentangling the ecological factors of selection, determining the timescales over which evolution occurs, and elucidating the architecture of such evolutionary patterns. Here, we employ experimental evolution to track the pace of the evolution of a common gut commensal, *Lactiplantibacillus plantarum*, within invertebrate (*Drosophila melanogaster*) and vertebrate (*Mus musculus*) hosts and their respective diets. While the symbiosis between Drosophila and *L. plantarum* is mainly determined by host diet, with the host influencing the fitness of its gut microbes only over short timescales, bacterial evolution within the mammalian host follows a divergent evolutionary path. Here, the host and its intrinsic factors play a critical role in selection and influence both the phenotypic and genomic evolution of its gut microbes, as well as the outcome of their symbiosis.

## INTRODUCTION

Millions of years of co-evolution between multicellular organisms and their microbial partners have resulted in mechanisms of mutual benefits consisting of complex networks of reciprocal interactions (Dethlefsen et al., 2007; Hooper et al., 2002; Ley et al., 2006; Moeller et al., 2016). One of the major melting pots of such a relationship is the gastrointestinal tract, where trillions of microorganisms form a rich and dynamic community collectively called the “gut microbiota”, which makes essential contributions to the host’s health (Backhed, 2005). In addition to aiding digestion (Hooper et al., 2002; Ley et al., 2008) and synthesizing essential metabolites (Gill et al., 2006; Nicholson et al., 2012), the gut microbiota is also involved in growth (Schwarzer et al., 2018; Storelli et al., 2011), organ development (Sjögren et al., 2012), immune system maturation (Bouskra et al., 2008; Hooper et al., 2012; Mazmanian et al., 2005), inflammatory responses (Garrett et al., 2010) and behavior (van de Wouw et al., 2017). Hence, the gut microbiota can be collectively thought of as a metabolically active organ integrated within the host (Bocci, 1992; O’Hara and Shanahan, 2006; Qin et al., 2010).

There is immense variation in the detail of the interactions between animals and their resident microbiota. In many insects, such as *Drosophila melanogaster*, the microbiome is reported to be fairly flexible, largely environmentally determined and less diverse than in mammals (Blum et al., 2013; Broderick et al., 2014; Chandler et al., 2011; Douglas, 2011; Early et al., 2017; Staubach et al., 2013; Storelli et al., 2018, 2011; Wong et al., 2014, 2013, 2011). On the contrary, mammals harbour trillions of microorganisms in their gut, which are known to stably colonize the gastrointestinal tract already during and after birth (Aagaard et al., 2014; Engel and Moran, 2013; Faith et al., 2013; Lee et al., 2013; Perez-Muñoz et al., 2017). Such assembly starts with low phylogenetic and species richness to culminate, over time, in the acquisition of a more complex and adult-like microbial profile (Bäckhed et al., 2015; Koenig et al., 2011; Palmer et al., 2007). Although gut microbes show higher resilience in mammals compared to many insects, the mammalian gut microbiota can also vary in response to both endogenous and environmental pressures (Candela et al., 2012; Flores et al., 2014; Hooper et al., 2002; Rawls et al., 2006; Walker et al., 2011). Indeed, the microbial ecosystem within the mammalian gut is shaped by the host’s genetic background (Benson, 2016; Falony et al., 2016; Goodrich et al., 2016, 2014; McKnite et al., 2012; Srinivas et al., 2013; Turnbaugh et al., 2009), together with anatomical, physiological, and immunological peculiarities (Rawls et al., 2006; Sousa et al., 2017). These include the intestinal architecture and composition (Johansson et al., 2008), the host’s innate and adaptive immune effectors (Salzman et al., 2010; Vaishnava et al., 2011; Macpherson et al., 2005), the host’s glandular secretions (i.e., gastric acid, bile, pancreatic fluids, and enzymes), as well as temperature and pH (Imhann et al., 2016; Jackson et al., 2016; O’May et al., 2005). At the same time, a plethora of environmental factors, largely related to the host’s dietary habits (depending not only on nutrient components, but also on the timing and regularity of consumption), and, in case of humans, the use of drugs (Cho et al., 2012; Forslund, 2015; Vich Vila et al., 2020; Zhernakova et al., 2016), the level of sanitization (Candela et al., 2012; Penders et al., 2006), practices related to infants’ delivery and feeding mode (Dominguez-Bello et al., 2011; Penders et al., 2006; Zivkovic et al., 2011), level of exercise (Karl et al., 2017), travel (David et al., 2014), and geographic location (Zhernakova et al., 2016), deeply contribute to the variation of such a microbial ecosystem.

In this light, several studies have sought to dissect the relative contributions of these factors in shaping the gut microbiota of insects and mammals. Among these, most have stressed the importance of the host’s diet as a key force in determining the microbiota configuration in both invertebrates (Erkosar et al., 2013; Jehrke et al., 2018; Obadia et al., 2018) and vertebrates (David et al., 2014; Hildebrandt et al., 2009; Muegge et al., 2011; Ridaura et al., 2013; Rothschild et al., 2018; Sonnenburg et al., 2016; Turnbaugh et al., 2009; Walker et al., 2011; Wu et al., 2011). In mice, switching from a low-fat, plant polysaccharide-rich diet to a high-fat/high-sugar “Western” diet can shift the microbiota structure within a single day (Turnbaugh et al., 2009), causing a progressive loss of species diversity over generations (David et al., 2014; Sonnenburg et al., 2016). Importantly, such diet-mediated microbiota alterations can ultimately result in specific microbiota–host layouts, which, in turn, affect host health and disease (Desai et al., 2016; Muegge et al., 2011).

Although these investigations have undoubtedly broadened our understanding of the diversity, resilience, and complexity of the gut microbiota across animal hosts, most of them have focused on characterizing how these external and internal factors shape microbiome compositional and functional features, eventually linking the resulting microbial pattern with a specific host trait (i.e., health or disease condition). Furthermore, by using common biomarkers (i.e., 16S rRNA), most of these studies have profiled the bacterial diversity at the genus or species level, therefore masking the potential presence of dynamic and rapidly evolving sub-populations. As a consequence, much less is known about the microbial evolutionary processes in the gut across animals. *In vivo* experiments, combined with deeper genome analyses, have recently brought forth a new appreciation of the gut microbes’ capability of rapidly diversifying and adapting in a newly colonized environment over short (Barreto, 2019; Barreto et al., 2020; Barroso-Batista et al., 2014) and long timescales (Yilmaz et al., 2021), providing insights into the ecological and molecular mechanisms underlying such evolutionary paths (Dapa et al., 2022, Fabich et al., 2011; Giraud et al., 2008; Lescat et al., 2017; Paepe et al., 2011; Ramiro et al., 2020). In this context, *Escherichia coli* is the most widely used model bacterium for studying bacterial evolution in the mammalian gut (Barreto et al., 2020; Barroso-Batista et al., 2014; Giraud, 2001; Giraud et al., 2008; Sousa et al., 2017). The reasons are manifold and linked to its ecological and clinical relevance, together with ease of experimental and genetic manipulation and tractability. However, further progress into understanding the drivers of microbial evolution in the gut of different animal hosts requires us to move beyond focusing on this particular species and to look at evolution in real time across a broader range of species. This is crucial for determining the main factors governing the evolution of gut microbes.

Here, we use *Lactiplantibacillus plantarum*, a common inhabitant of the gastrointestinal tract of different animals (Marco et al., 2009), as model species to explore the evolutionary trajectories of gut microbes across animal hosts, and particularly if and how they differ between insects and mammals.

By using *Drosophila melanogaster* as an animal model, we previously demonstrated that the host’s diet, rather than the host environment *per se*, is the predominant force in driving the emergence of the symbiosis between *L. plantarum* and the fruit fly (Martino et al., 2018). Here, we hypothesize that, given the higher persistence and colonization ability of gut microbes in the gastrointestinal tract of mammals, the mammalian host and its intrinsic factors represent key agents of selection in the evolution of gut microbiota as compared to the mammalian host’s diet. To explore this, we performed a parallel experimental evolution of the bacterial strain *L. plantarum* NIZO2877, which was previously shown to moderately promote growth both in *Drosophila* and mice (Schwarzer et al., 2016). We evolved *L. plantarum* in mono-association with germ-free C57Bl6 mice and in the mouse laboratory diet, separately. At the same time, we experimentally evolved the same strain in association with *Drosophila* and its nutritional environment to test if and how the presence of the invertebrate host affects the tempo and mode of evolution of its gut microbiota.

Our results indicate that the evolution of the same gut bacterium diverges between insects and mammals, pointing out the effects of the different host-derived selection pressures. While, in *Drosophila*, the host seems to benefit the fitness of *L. plantarum* in a short timescale, without significantly affecting its evolutionary trajectory, microbiome evolution in mammals follows a completely different path. Here, host factors represent a crucial agent of selection, shaping the evolution of gut microbes both on a phenotypic and genomic level. This results in an increased bacterial adaptation toward the host’s intestinal environment, revealing new insights into the symbiosis between *L. plantarum* and mammalian hosts.

## RESULTS

### *L. plantarum* evolution within a mammalian host leads to higher adaptation to the host intestinal environment

With the aim of investigating the evolutionary dynamics of *L. plantarum* in a mammalian host and to dissect the role of the mammalian host’s diet in the evolution of gut microbes, we experimentally evolved the bacterial strain *L. plantarum*^NIZO2877^ (*Lp*^NIZO2877^) in the intestine of germ-free (GF) C57Bl6 mice (Host setup) and in the mouse laboratory diet (Diet setup) separately (Figure 1a), following the same experimental setup that we have previously applied to *Lp* evolution in *Drosophila* (Martino et al., 2018). Specifically, in the Diet setup, we mono-associated the mouse diet with the ancestral strain in the absence of the host, and monitored microbial evolution for 20 transfers (T) in five independent replicates (i.e., 20 days, corresponding to ∼400 bacterial generations). In the Host setup, we mono-colonized 7 GF mice housed in cages in one Trexler-type isolator with the ancestral strain by intragastric gavage; once the mono-association had been performed, 4 female mice were bred to generate F0 generation for the next 10 months and 2 females and 1 male were the F0 founders of the subsequent mouse generation. Bacterial evolution was followed across F0 and the subsequent generations of mice (F1, F2, F3, and F4) for 10 months (i.e., ∼286 *Lp* bacterial generations; Figure 1a,b). Evolving bacteria were horizontally dispersed and vertically transmitted across generations with no further artificial inoculation. Over this period, the mice were fed with the same diet used for the Diet setup.

**Figure 1.**
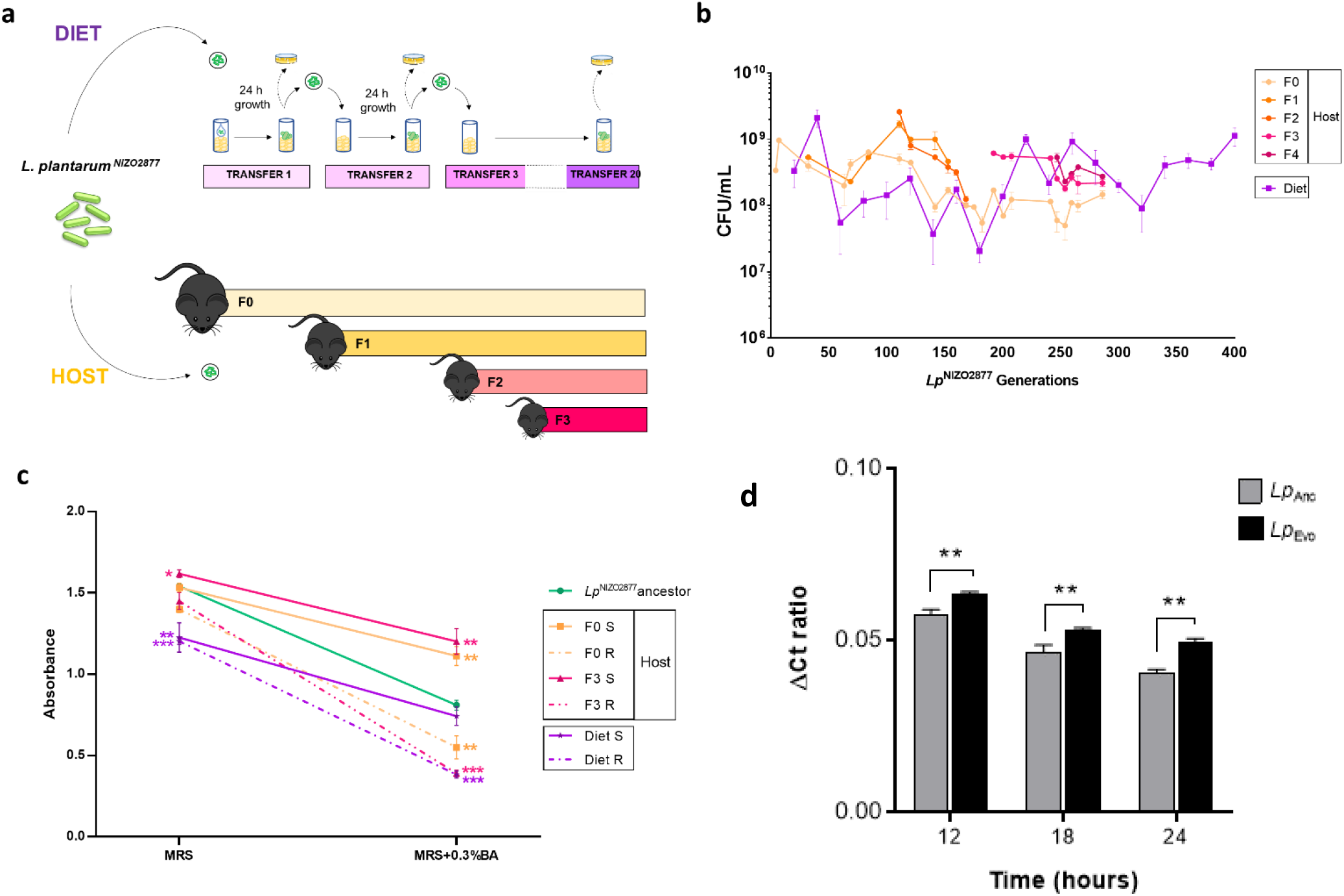
(a) Schematic representation of the *Lp*^NIZO2877^experimental evolution (EE) protocols in the mouse diet (Diet setup) and mouse intestine (Host setup). The ancestral strain was inoculated in the mouse laboratory diet and in the mouse intestine by intra-gastric gavage. Five and two replicates of EE were performed for the Diet and the Host setup respectively. An evolved bacterial subpopulation was plated out on MRS agar plates for colony counting at different time points throughout the EE in both setups. The EE in the Diet setup was conducted up to 20 transfers (i.e., ∼20 days, corresponding to approximately 400 *Lp* generations). In the mouse intestine, *Lp* EE was carried out from generation 0 (F0) to 4 (F4) of mice (i.e., 10 months after the mono-association, ∼300 days, corresponding to ∼286 Lp generations) for one replicate and for one generation (F0) (3 months after the mono-association) for the second replicate. (b) *Lp*^NIZO2877^ growth trend monitored over the course of the EE in both experimental setups (Host and Diet). (c) Final absorbance values (OD = 600) reached by the *Lp* strains under standard growth conditions (MRS broth) and in MRS broth added to with 0.3% bile acids (BA). S indicates smooth colonies, while R indicates rough colonies. Each dot represents the mean of at least three experimental replicates per condition, with bars indicating the respective SD (standard deviation). For both experimental conditions (MRS broth and MRS broth + 0.3% BA), asterisks indicate significance between the final absorbance values of each strain against those of the *Lp*^NIZO2877^ancestor (unpaired t-test; *p ≤ 0.05, ** p < 0.01, and *** p < 0.001). (d) Quantitative PCR analysis of *Lp*^NIZO2877^ ancestral strain (Anc) and *Lp*^NIZO2877^-derived population (Evo) evolved in the mouse intestine. Lines above each bar represent the mean with the standard deviation (SD) of normalized ΔC_T_ ratios (1/ΔC_T_) of 3-4 mice/group, Statistical significance of the results is included (Student’s t test, *p≤0.05, **p≤0.01).

Overall, *L. plantarum* growth showed a similar trend across the two evolutionary setups. In the mouse diet, we found that the microbial load increased extremely fast, as it shifted from the initial inoculum of 10^2^ total CFU/mL to 3.34 × 10^8^ CFU/mL in less than 24 h (Figure 1b). Such a growth trend was maintained over the course of the evolution experiment, during which the bacterial load ranged between 10^7^ and 10^9^ CFU/mL, with the lowest value at T9 (i.e., after ∼180 *Lp* generations, Mean_T9_ = 2.06 × 10^7^ CFU/mL) and the highest at T2 (i.e., ∼40 *Lp* generations, Mean_T2_ = 2.12 × 10^9^ CFU/mL) (Figure 1b). Microbial growth within the mouse intestine showed comparable loads as the Diet-evolved populations, reaching the highest peak after ∼110 *Lp* generations (F2 = 2,6 × 10^9^ CFU/g feces) and the lowest after ∼260 bacterial generations (F0, Mean = 5 × 10^7^ CFU/g feces). Moreover, within each mouse generation, no significant differences were detected when comparing the initial and final bacterial concentrations (Figure 1b). These results suggest that *L. plantarum* was able to reach and maintain high abundance both in the mouse diet and in the mouse intestine.

To compare *L. plantarum* phenotypic evolution with and without the mammalian host, we conducted a morphological analysis of the *Lp* evolved colonies over time in both experimental setups. Interestingly, we detected the appearance of an evolved *L. plantarum* sub-population showing a different colony morphology compared to the ancestral strain in both evolutionary setups. Specifically, while *L. plantarum* typically forms rounded, smooth colonies on MRS agar, the newly evolved colonies showed a less-defined, rough morphology, which looked more transparent than the ancestral one (Figure supplement 1a). Such a population appeared after ∼57 *Lp* generations in the Host setup (F0) and after ∼280 *Lp* generations in the Diet setup (T14). In the mouse intestine, it initially affected 48% of the whole bacterial population, but it tended to decrease over time, reaching 38% of the population during mice Generation 3 (*i*.*e*., *∼*140 *Lp* generations, Figure supplement 1d). To further characterize *L. plantarum* phenotypic change, we analyzed the two bacterial morphotypes with an electron microscope. Interestingly, while the ancestral-like colonies showed a compact bacillary structure with a smooth surface, the newly evolved rough colonies had an irregular surface, and looked elongated and filamentous (Figure supplements 1b-c). However, it was unclear whether such longer chains resulted from undivided bacterial cells, or whether they corresponded to single cells. Notably, when a rough colony was re-streaked on MRS agar or cultured in MRS broth, the morphology reversed to a smooth, ancestral-like one, showing that the newly evolved phenotype was transient and reversible (data not shown).

We next sought to investigate the mechanisms underlying the emergence of the new microbial morphotype. Bacterial morphological changes commonly occur as a result of stress responses to a wide range of factors (Bron et al., 2004; Gandhi and Shah, 2016; Ingham et al., 2008; Yang et al., 2016). In the mammalian intestine, one of the most stressful conditions is due to the activity of bile acids (BA) (Bron et al., 2004; Hussain et al., 2013; Merritt and Donaldson, 2009; Sistrunk et al., 2016; Urdaneta and Casadesús, 2017). We thus hypothesized that the presence of bile acids encountered during transit through the mouse gastrointestinal tract might have contributed to the appearance of the newly evolved rough morphology. On the contrary, the appearance of the rough morphology in the Diet-evolved populations could not have resulted from a stress response to bile acids, since they are absent in the mouse diet. As a consequence, we expected that both Diet-evolved colonies (rough and smooth morphotypes) would be equally affected by the presence of bile acids.

To verify these hypotheses, we first tested whether bile acids are able to induce a stress response in *L. plantarum*. We serially propagated the ancestral strain *Lp*^NIZO2877^ in MRS broth and MRS broth with the addition of 0.3% BA, corresponding to the physiological concentration found in the mouse intestine (Stenman et al., 2013) for seven days and measured microbial growth. As expected, bacterial loads were significantly lower in presence of 0.3% BA compared to the control (MRS) already after one Transfer (*Lp*_T1-MRS+0.3%BA_ = 4.07 × 10^6^ CFUs; *Lp*_T1-MRS_ = 1.75 × 10^8^ CFUs) throughout the experiment (*Lp*_T7-MRS+0.3%BA_ = 8.06 × 10^7^ CFUs; *Lp*_T7-MRS_ = 3.14 × 10^11^ CFUs) (Figure supplement 2). By monitoring the bacterial morphology on agar plates, we noticed that, although the *Lp* colonies grown in the presence of BA looked slightly smaller and more transparent, the colonies maintained the ancestral smooth morphotype over the seven serial transfers (data not shown). These results demonstrate that bile acids are able to induce a stress response in *Lp*^NIZO2877^ by impairing bacterial growth.

Next, we analyzed the fitness of Diet- and Host-evolved bacteria in the presence of bile acids and detected a *Lp-*specific response to BA depending on the morphology and evolutionary background. In detail, all rough colonies reached the lowest final absorbance values in the presence of bile acids, regardless of their evolutionary history (i.e., Host- or Diet-evolved) (Figure 1d, Figure supplements 3 and 4). They also showed a slower and delayed growth in standard MRS broth compared to the ancestor (Figure supplement 3). Remarkably, while the Diet-evolved smooth morphotypes reached a significantly lower absorbance value compared to the ancestor (Figure supplement 3), we detected signatures of adaptation toward bile acids among the Host-evolved smooth colonies (F0, F3). Such colonies reached, overall, the highest absorbance values in presence of bile acids (Figure 1c), suggesting higher tolerance to the stress compound.

To investigate whether *L. plantarum* populations evolved in the mouse intestine exhibited signs of adaptation to the animal host, mice bearing a conventional gut microbiota were gavaged either with the *Lp*^NIZO2877^ ancestral strain or with the *Lp*^NIZO2877^-derived population evolved in the mouse intestine for ten months (sample F0-10). Whole bacterial DNA was isolated from feces and real-time PCR was performed to track *L. plantarum* persistence over time (i.e., up to 72 h). Notably, we observed a significantly higher persistence of the *L. plantarum*-evolved population compared to the ancestral one until 24 h after gavage, while no *L. plantarum* was detected after 36 h (Figure 1d).

Altogether, our data demonstrate that *L. plantarum* populations evolved in the mouse intestine showed adaptation towards the animal host. This, among other factors, may result from the increased tolerance to intestinal stress (i.e., presence of bile acids), and leads to higher and longer persistence in the host intestinal environment.

### Bacterial evolution within the mammalian host intestine is shaped by the emergence of hypermutators and a higher number of mutations compared to the evolution in the host diet

To investigate the influence of the mammalian animal host and its nutritional environment on the genomic evolution of its symbiotic bacteria, we sequenced the genomes of whole bacterial populations evolved in the mouse diet and in the mouse intestine at different time points. Specifically, we sequenced the genomes of fifteen *L. plantarum* evolved populations isolated from faeces pooled from 3-4 individual mice across the five generations from one replicate of experimental evolution (EE) (Supplementary file 1). As for the Diet setup, we sequenced the genomes of six evolved bacterial populations isolated during Transfers 2 and 14 from three independent replicates out of five (Supplementary file 1). In addition, we sequenced the genomes of single bacterial colonies showing the smooth, ancestral-like phenotype and the newly evolved rough morphology (replicate 2, Transfer 14, Supplementary file 1).

Bacteria propagated in the mouse diet showed overall a low number of mutations, with the highest value in T14 (N= 8 mutations) (Table 1, Figure 2a, Supplementary file 2).

**Table 1.**
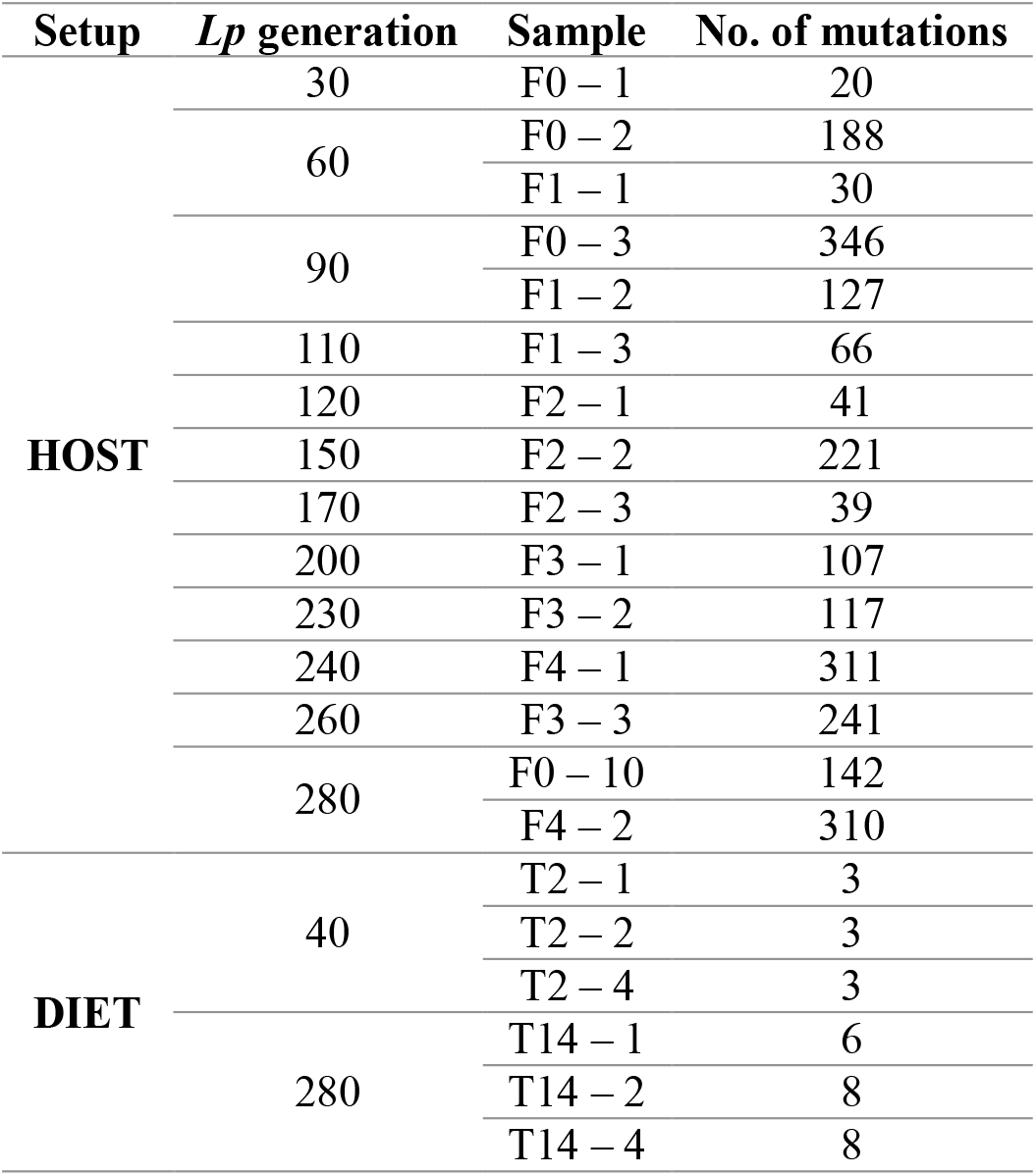
Number of mutations detected for each *L. plantarum* evolved population in the Host setup (F0, F1, F2, F3, and F4 generations) and the Diet setup (Transfers 2 and 14).

**Figure 2.**
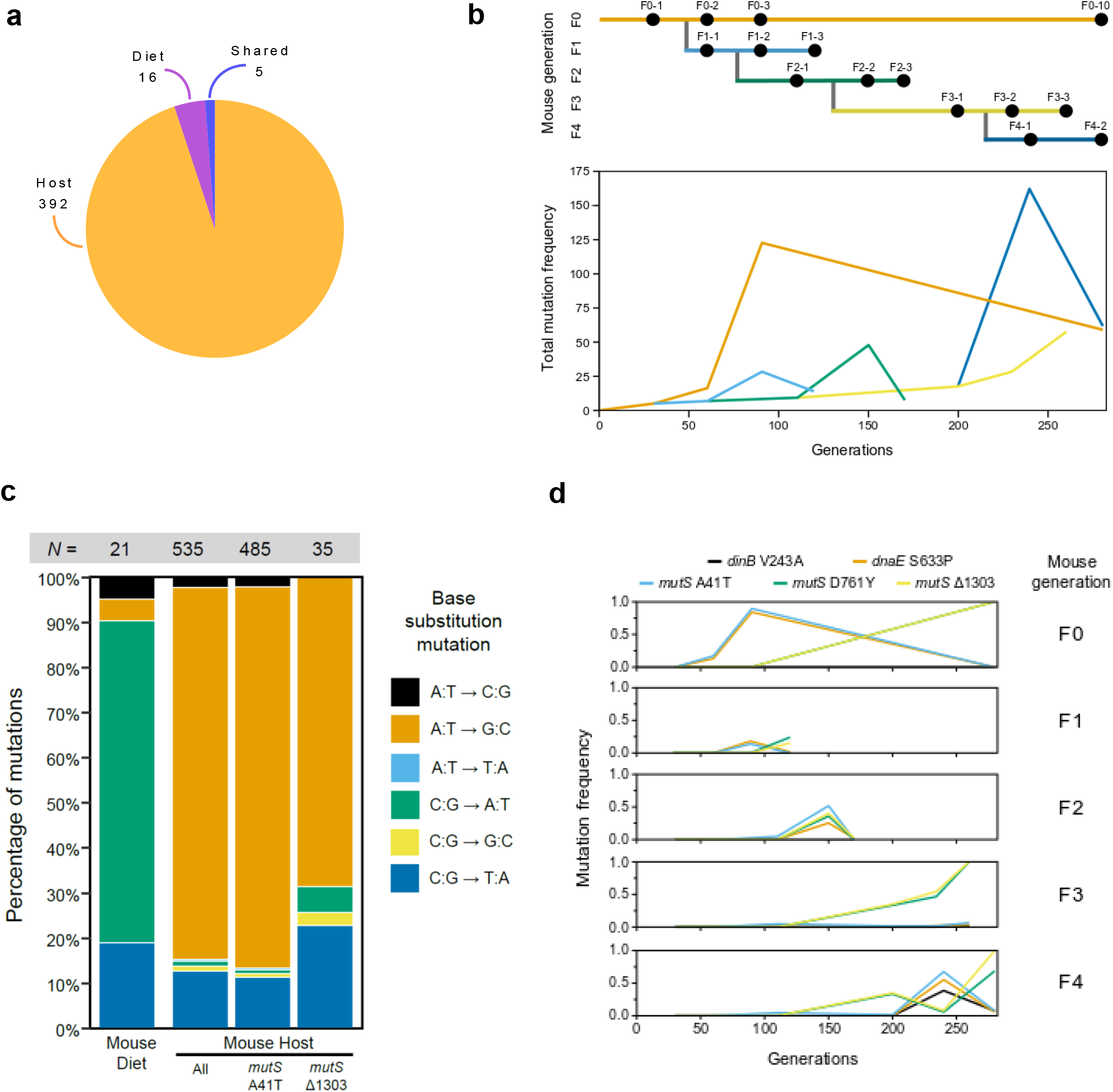
Mutations that occurred during *L. plantarum* experimental evolution in mice. (a) Pie-chart reporting the number of *Lp* genes that accumulated mutations over the course of each experimental evolution setup (Host- and Diet-exclusive mutations) and in both setups (Shared). (b) Total summed frequencies of all mutations observed in each sample from the mouse host evolution experiment. The panel above shows the structure of the evolution experiment in mouse hosts. Bacterial populations were sequenced at the indicated time points. (c) Base substitution spectra observed in the mouse diet, as well as mouse host populations divided into all mutations and those associated with the *mutS* A41T and *mutS* Δ1303 mutator lineages (see Figure Supplement 5). The total number of mutations detected in each group (*N*) is shown above each bar. (d) Frequency trajectories of the *mutS* A41T and *mutS* Δ1303 mutations and other mutations in DNA replication and repair genes (*dinB, dnaE*) that may alter mutation rates in the mouse host populations.

During this time point, the single smooth and rough colonies showed the emergence of only 1 and 3 mutations per strain, respectively (Replicate 2, Supplementary file 3). On the contrary, bacteria evolved in the mouse intestine revealed a marked increase in the total number of mutations compared to the Diet-evolved ones (Table 1, Figure 2a,b, Supplementary file 4). Remarkably, the high number of genetic variants detected in the Host-evolved *Lp* populations correlated with the appearance of novel mutations in the genes encoding for DNA mismatch repair proteins *mutS* (methyl-directed mismatch repair complex subunit), *dnaE* (DNA polymerase III alpha subunit), and *dinB* (DNA polymerase IV) (Supplementary file 4).

Mutations in the mouse host populations were overwhelmingly A:T to G:C base pair substitutions, whereas these substitutions accounted for less than 10% of mutations in the mouse diet (Figure 2c). An elevated A:T to G:C mutational bias is consistent with mutations in *mutS* (Schaaper and Dunn, 1987) but not *dinB* (Wagner and Nohmi, 2000) or *dnaE* (Mo et al., 1991), so we concluded that the evolved *mutS* alleles are probably largely responsible for the hypermutator phenotype. To test whether hypermutation of *L. plantarum* in mouse was repeatable, we Sanger sequenced the *mutS* gene of the *Lp* populations evolved in the second replicate of experimental evolution in the mouse intestine (F0 – sequenced time points: 1, 2 months). Here, we detected five additional variants in the *mutS* gene, suggesting the appearance of independent hypermutator lineages (Supplementary file 5).

Whole genome sequencing of the first replicate of *Lp* EE in mice detected three separate mutations in the *mutS* gene (Supplementary file 4). A point mutation causes an A41T amino acid substitution. A deletion of base pair 1303 of the *mutS* coding sequence results in a frameshift. Another point mutation would lead to a D761Y amino acid substitution. The frequency of this last mutation tracked with that of the Δ1303 mutation as soon as they both reached observable frequencies (Figure 2d). Since the D761Y mutation is located after a new stop codon in the *mutS* gene created by the Δ1303 mutation at codon 456, it would not affect the *mutS* gene product in the context of this mutation. Therefore, we considered the A41T and Δ1303 mutations as defining two distinct *mutS* hypermutator lineages.

The two *mutS* hypermutator lineages and a nonmutator lineage that maintained the ancestral mutation rate competed throughout the history of the evolution experiment in different animals (Figure 2d). The *mutS* A41T lineage appeared first and increased in frequency over the initial three months (63 generations) in the F0 mouse, reaching up to a 90% frequency. Between the third month (F0-3) and tenth month (F0-10) however this lineage fell below the level of detection and the *mutS* Δ1303 lineage rose to near 100% frequency. In the F3 and F4 mouse generations the *mutS* Δ1303 lineage also ultimately swept to 100% frequency. In the F1 mouse however neither lineage exceeded 20% frequency while, in the F2 mouse generation, both *mutS* lineages increased to ∼40-50% frequency between generations 82 (F2-1) and 104 (F2-2) but then declined to less than 3% frequency by generation 123 (F2-3).

By fitting a model that assumed both *mutS* alleles evolved near the beginning of the experiment and accounted for a certain fraction of the population at each time point, we were able to estimate that the *mutS* A41T lineage accumulated mutations at a rate of 0.980 ± 0.092 per generation and the *mutS* Δ1303 lineage accumulated mutations at a rate of 0.170 ± 0.038 per generation (± values are standard errors of fit values). These rates are 142- and 25-fold the rate in the mouse diet treatment. The roughly 6-fold factor by which the two *mutS* lineage rates differ from one another is significant (F-test, *p* = 5.3 × 10^−6^). It may indicate that the *mutS* alleles have different effects on their own or in combination with mutations in other genes. For example, the *dnaE* mutation tracks with the *mutS* A41T allele throughout its entire history, and the *dinB* mutation appeared in this genetic background later in the F4 mouse (Figure 2d). By comparing the genetic variants detected in the Host- and Diet-evolved populations, we identified 5 genes that mutated in both evolutionary setups (Figure 2a, Supplementary files 2,4). However, none of them persisted across generations of both conditions. We then turned our attention to those targets that exclusively mutated in the Host and Diet setups and that showed persistence across EE cycles. While within the Diet-evolved populations, we only detected two mutations (i.e., genes: dipeptidase, hypothetical protein; Supplementary file 2), the *Lp* populations evolved in the mouse intestine showed a significantly higher number of mutations (*n*= 310; Supplementary file 4), which were detected in 41 genes (and six intergenic regions). To understand whether such variants were linked to specific functions, we clustered them according to their predicted functional category. While seven of the 41 mutated genes were related to unknown functions, 30 genomic targets were grouped in 12 total functional categories, among which the carbohydrate transport and metabolism was the most enriched category (Figure 3 and Supplementary file 6). Notably, four mutational targets belonging to this functional category included genes encoding for phosphotransferases system (PTS) transporters (*PTS*—*mannose-specific IIC, PTS— sucrose specific IIA/IIB/IIC, phosphocarrier protein of PTS system*, and *PTS system—beta-glucoside-specific IIB/IIC/IIA component*). In addition, the genes belonging to the signal transduction and metabolism (*n* = 5) and transcription (*n* = 5) categories resulted to be affected by mutations in the Host-evolved populations (Figure 3 and Supplementary file 6).

**Figure 3.**
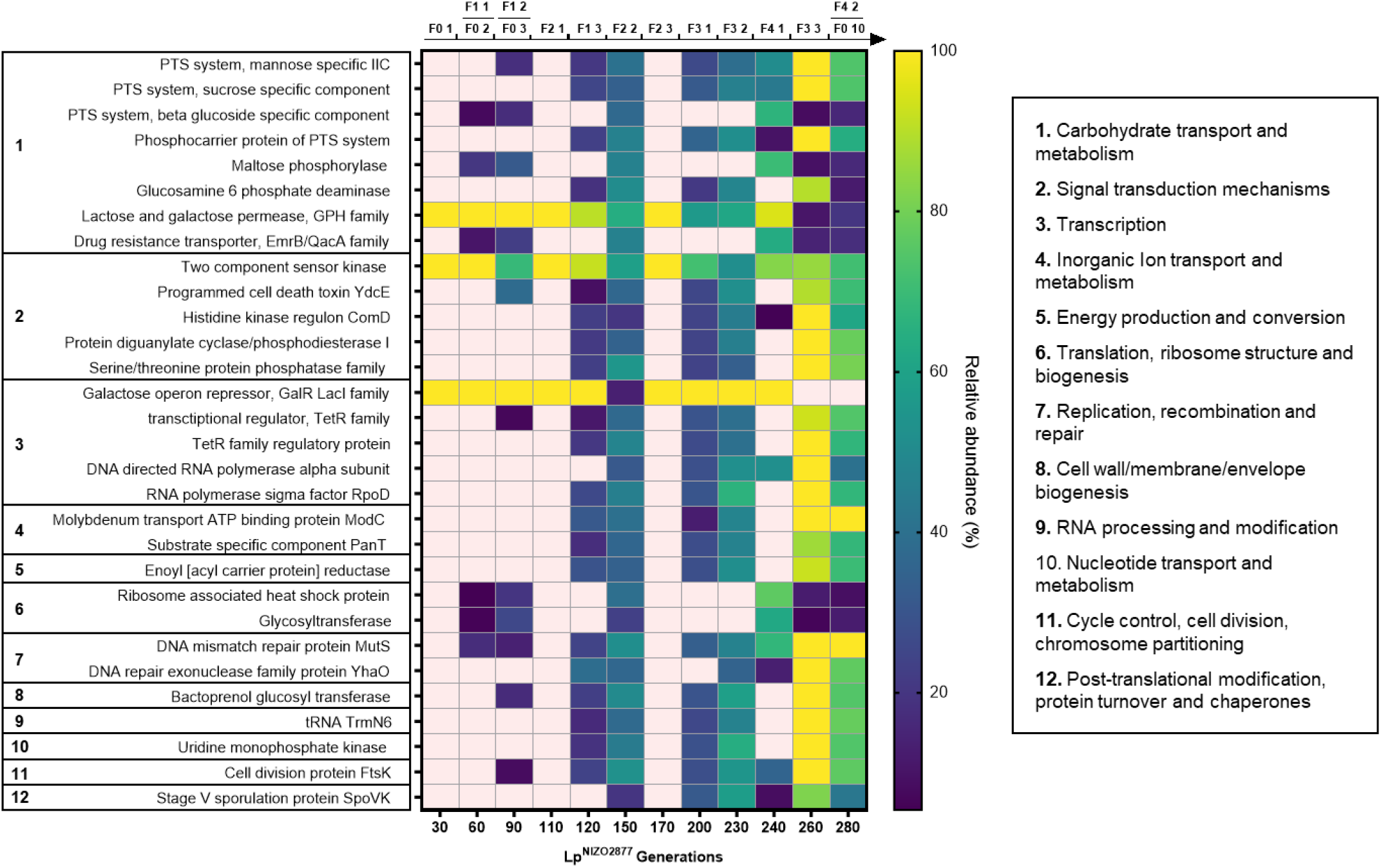
*L. plantarum* genes with mutations that persisted over the course of the evolution experiment in the mouse intestine. Each target is grouped according to the predicted functional category. The colors of the heatmap indicate the relative abundance of each mutational target across time points. Lighter spaces indicate that no mutations were detected. The timeline above the heatmap represents the samples from which the each *Lp* population was retrieved and sequenced.

Altogether, our results demonstrate that *L. plantarum* follows divergent evolutionary paths in the mouse intestine compared to the mouse diet, both genomically and phenotypically. While *L. plantarum* evolution in the mouse diet was characterized by low mutational load, hypermutators were detected during evolution in the mouse intestine. In addition, *L. plantarum* populations evolved inside the host exhibited higher adaptation to host intrinsic factors (i.e., presence of bile salts).

### *Drosophila melanogaster* benefits *L. plantarum* growth on a short time scale

The ecological and evolutionary dynamics of gut microbes vary greatly between mammals and insects. Among many factors, this is largely due to the different extent through which microbes colonize the gut of animals and, as a consequence, the different degree of dependence between animals and microbes across hosts (Blum et al., 2013; Broderick et al., 2014; Cen et al., 2020; Chandler et al., 2011; Douglas, 2011; Early et al., 2017; Marco et al., 2009; Moran et al., 2019; Staubach et al., 2013; Storelli et al., 2018, 2011; Wong et al., 2011). By using *Drosophila melanogaster* as animal model, we have previously demonstrated that, in *Drosophila*/*L. plantarum* symbiosis, the host nutritional environment, rather than the host *per se*, is the predominant force in driving the emergence of such symbiosis (Martino et al., 2018). The experimental setup was comparable to the one used in the present study to experimentally evolve *L. plantarum* in the mouse intestine and in the mouse diet. Specifically, in the Host setup, bacteria were horizontally and vertically transmitted among individuals and across generations, while in the Diet setup, artificial passages of evolving bacterial populations were conducted through experimental generations (Martino et al., 2018). We then asked whether such differences in the selection regime between setups (natural transmission of bacteria in the Host setup *versus* artificial inoculation in the Diet setup) might have influenced the microbial evolutionary dynamics both in the fruit fly and in mice. To test this, we decided to replicate the *Lp* experimental evolution with and without *Drosophila* (Host and Diet setup respectively) for a total of 20 cycles (220 days, corresponding to ∼1760 bacterial generations) by applying the same transfer and sampling time for both setups (five independent replicates per setup, Figure 4a). In this way, we were able to minimize differences between setups so that the microbial evolutionary trajectories relied exclusively on the presence/absence of the animal host.

**Figure 4.**
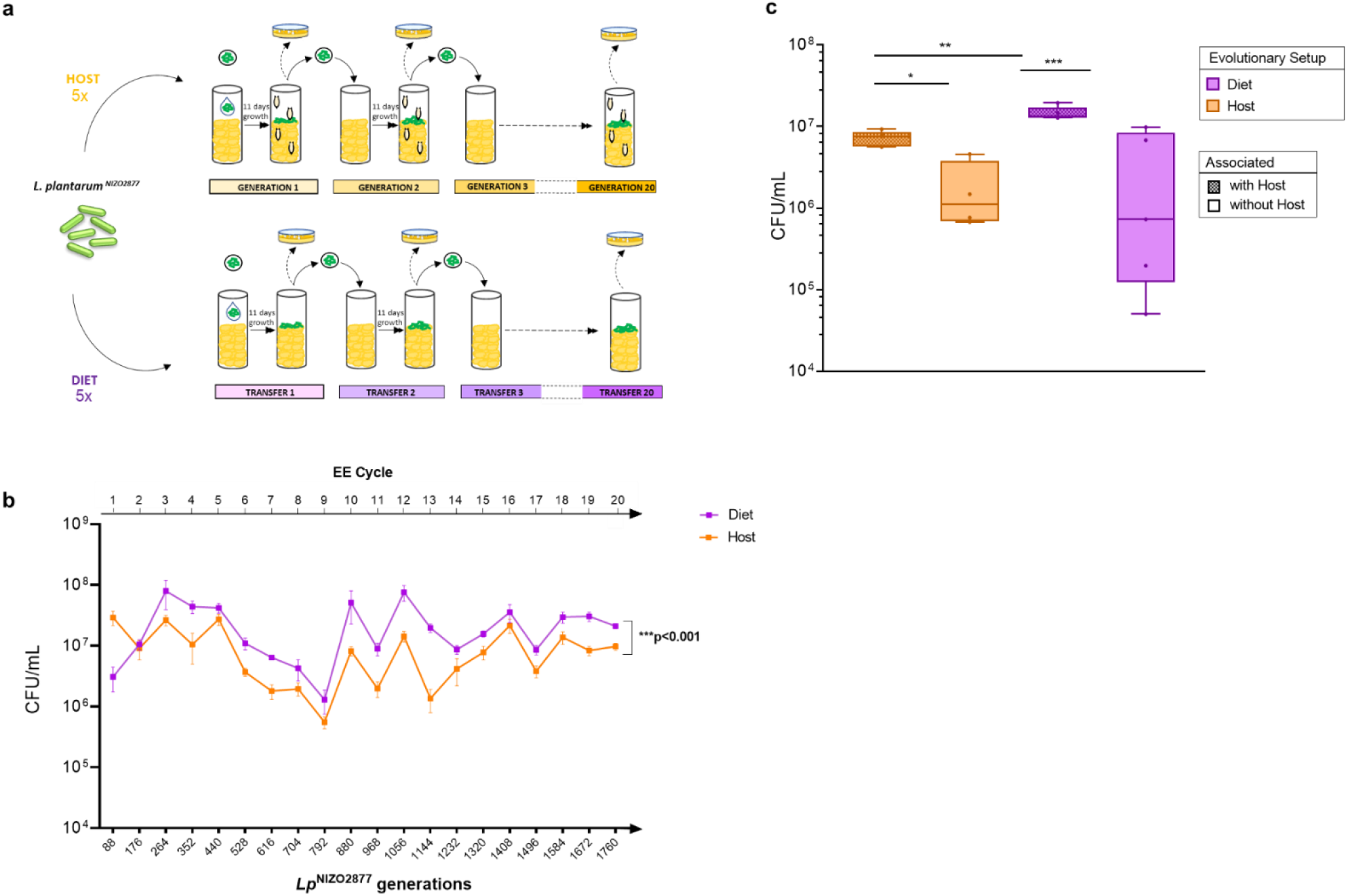
(a) Design of the *Lp* experimental evolution (EE) with and without Drosophila (Host and Diet setups, respectively). For the first EE cycle, the ancestor strain (*Lp*^NIZO2877^) was inoculated into tubes containing a poor-nutrient Drosophila diet (Diet setup) or a poor-nutrient diet containing 40 germ-free Drosophila embryos (Host setup). No further inoculation of the ancestor was performed until the end of the experimental evolution. As soon as at least 15 pupae emerged from all host tubes (i.e., after ∼11 days, corresponding to ∼88 bacterial generations), 150 µl of food were collected from both setups using a sterile loop, homogenized, and plated out to isolate bacteria (frozen “fossil” records of EE cycle 1). This bacterial population was used as the inoculum for the following generation/transfer. Subsequent EE cycles followed the same experimental procedure as cycle 1 and started from the fossil records belonging to the previous generation/transfer. The EE lasted 20 cycles (220 days, corresponding to ∼1760 bacterial generations). (b) *Lp* growth over the course of the Host and Diet EE protocols across 20 total EE cycles (i.e., 1760 bacterial generations). Each point represents the mean of the five experimental replicates, with bars indicating the standard error of the mean (SEM). ANCOVA *** p < 0.0001). (c). Microbial load obtained by mono-associating each of the five replicates of both setups (Host- and Diet-evolved bacteria) isolated from cycle 17, both with and without the host. Each bar represents the standard error of the mean (SEM) obtained by analyzing five replicate populations for each condition. Ordinary one-way ANOVA (* p ≤ 0.05, ** p <0.01, and *** p < 0.001).

We first investigated whether the presence of *Drosophila* affected *Lp*^NIZO2877^ evolution by conducting morphological evaluations of the evolved bacterial populations and analysing the microbial growth dynamics throughout the two experimental protocols. Specifically, the evolved bacterial populations in the ten independent replicates have been plated out at the end of each experimental cycle (total of 20 cycles), macromorphological evaluation of bacterial colonies was routinely performed and microbial load was measured at the end of each cycle (∼11 days). Contrary to what we observed during *L. plantarum* evolution in the mouse intestine, no differences in colony morphology were observed across the *Lp* populations evolved with and without *Drosophila*. Remarkably, the microbial load was significantly higher overall in the absence of the host, except for in the first generation (i.e., 88 bacterial generations) (Figure 4b). Since the difference in microbial growth between the two setups was detected at a specific time point (i.e., the end of each experimental generation - 11 days), we asked whether the higher microbial load observed in the Diet setup resulted from a delayed growth dynamic compared to that of the Host-evolved populations. To address this question, we analyzed the microbial concentration at an earlier time point (i.e., seven days after the mono-association). The *Lp* concentration was confirmed to be overall significantly higher in the absence of the host, except for generations 6, 19, and 20 (i.e., after ∼528, 1672, and 1760 bacterial generations, respectively) (Figure supplement 5a).

Interestingly, throughout our sampling period, we also noticed an unexpected correlation between the growth dynamics of the Host- and Diet-evolved populations, which was particularly pronounced from generation 6 to 12 (i.e., from 528 to 1056 *Lp*^NIZO2877^ generations) (Figure 4b). To test the repeatability of our findings, we replayed *L. plantarum* EE from cycles 5 to 12 in both setups and analyzed the microbial concentration after 7 and 11 days. Our results further confirmed the correlation in microbial growth dynamics between the Host and Diet setup (Figure supplement 5). This demonstrates that *Lp* growth dynamics are repeatable and did not result from experimental artefacts or external variables.

We then asked whether and how *Lp* evolutionary history (i.e., evolving in presence or absence of its host) could generate evolutionary trade offs in a different environment. To this end, we mono-associated *Lp* Host and Diet-evolved populations isolated at the end of EE cycle 17 from each of the ten independent replicates (five replicates per setup), both in the presence and absence of *Drosophila* and analyzed the microbial load in both conditions after 11 days. *Lp* concentration was always significantly higher in the presence of *Drosophila* (Figure 4c). This was also visible after cycle 1 of *Lp* EE (i.e., ∼88 *Lp* generations; Figure 4b). However, when comparing the microbial evolutionary backgrounds, the Diet-evolved populations reached significantly higher loads compared to the Host-evolved populations when associated with the fruit fly (Figure 4c). Taken together, our results show that, although *Drosophila* benefits *L. plantarum* growth on a short timescale, bacterial evolution ultimately leads *L. plantarum* to grow to a higher extent in the absence of *Drosophila*.

### Genome sequencing reveals parallel genomic evolution between *Lp* populations evolved with and without *Drosophila*

We next investigated if and how *Drosophila* influences *Lp* evolution on a genomic level. To do this, we performed metagenomic sequencing of bacterial populations isolated from both experimental setups during EE cycle 2, 8, 14 and 20 (three independent replicates sequenced per time point and setup, Supplementary material 1). Across the two evolutionary setups, we detected a similar mutational trend in terms of the number of genetic variants (Table 2, Figure 5a,b) and identified signatures of strong parallel genomic evolution. Bacteria evolved in the fly host populations accumulated 0.00177 ± 0.00018 mutations per generation and fly diet populations accumulated 0.00150 ± 0.00010 mutations per generation (± values are standard errors of fit values). These rates did not differ significantly from one another (*F*-test, *p* = 0.059). Mutations in the fly host and diet populations were mainly C:G to A:T and C:G to T:A substitutions (Figure 5c).

**Table 2.** Number of mutations detected for each *L. plantarum* evolved population in the Fly Host and Diet setup.

| EE cycle | Sample | No. of mutations<br>HOST setup | No. of mutations<br>DIET setup |
| --- | --- | --- | --- |
| 2 | 1 | 2 | 4 |
|  | 2 | 8 | 3 |
|  | 4 | 4 | 2 |
| 8 | 1 | 10 | 6 |
|  | 2 | 8 | 8 |
|  | 4 | 7 | 3 |
| 14 | 1 | 7 | 9 |
|  | 2 | 7 | 8 |
|  | 4 | 7 | 4 |
| 20 | 1 | 7 | 5 |
|  | 2 | 6 | 8 |
|  | 4 | 6 | 3 |

**Figure 5.**
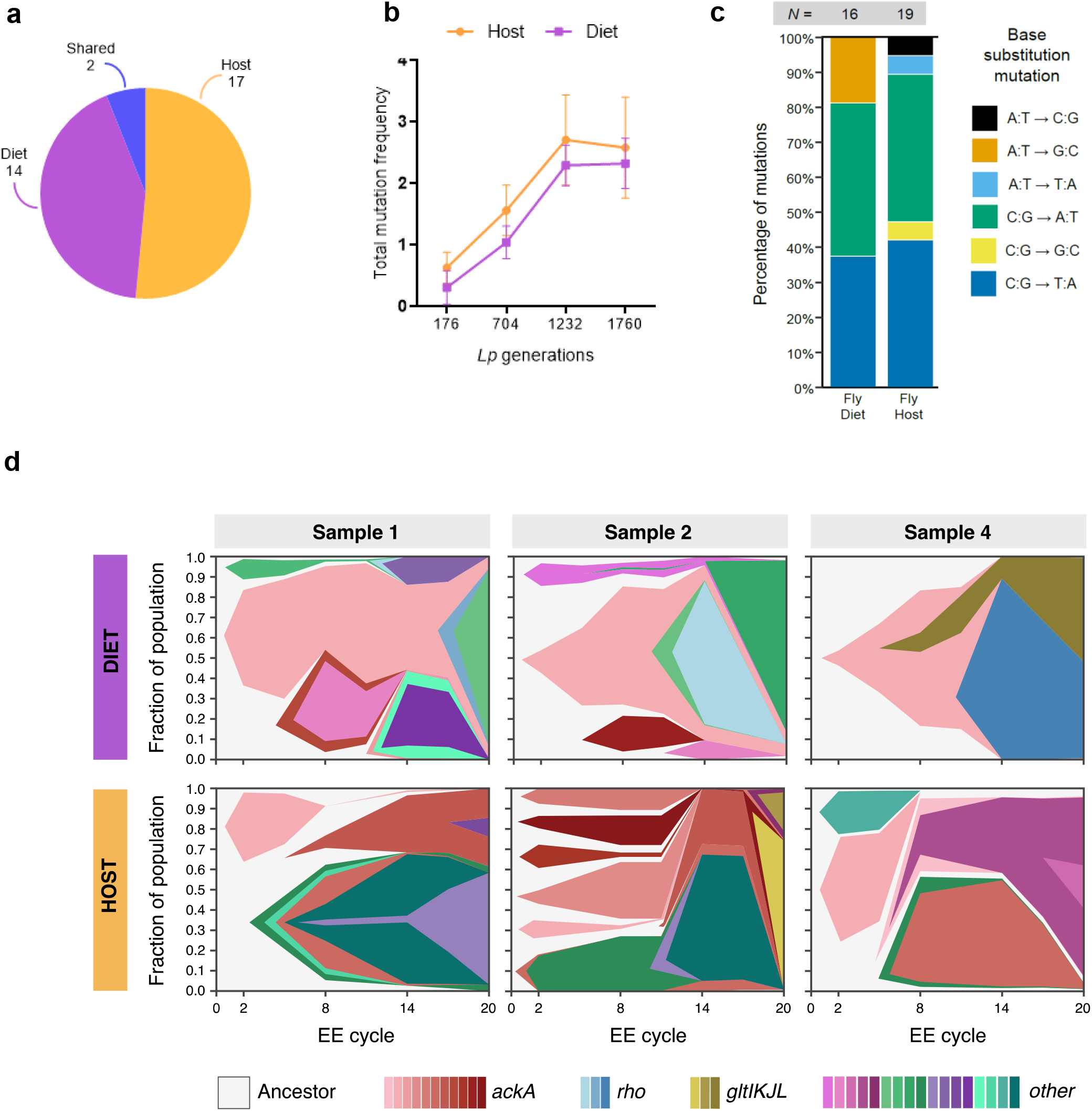
Mutations occurred during *L. plantarum* experimental evolution in the fly setup. (a) Pie-chart reporting the number of *Lp* genes that accumulated mutations over the course of each experimental evolution setup (Host- and Diet-exclusive mutations) and in both setups (Shared). (b) Total summed frequencies of all mutations observed in each sample from the Host and Diet evolution experiment. Each point represents the mean of the 3 experimental replicates analysed, with bars indicating the standard deviation (SD). (c) Base substitution spectra observed in the fly host and diet. The total number of mutations detected in each group (*N*) is shown above each bar. (d) Dynamics of mutations in fly diet and host treatments. Muller plots representing the evolutionary dynamics of *Lp* genes that mutated in the Host and Diet-evolved populations across EE cycles 2, 8, 14 and 20 (i.e., after 176, 704, 1232 and 1760 *Lp* generations).

Two genes mutated in both setups (i.e., *ackA* and *gltL*, Figure 5c and Supplementary files 7, 8). Specifically, all *Lp* experimental replicates isolated from the Host and Diet setups showed at least one genomic change within the acetate kinase A gene (*ackA*), which was the first gene to be affected by mutation and exhibited the highest number of mutations per gene (Figure 5c, Supplementary files 7, 8). This result confirms our previous work showing that mutations of *L. plantarum ackA* occur both in the presence and in absence of *Drosophila*. We identified 12 non-synonymous *ackA* mutations, one of which was shared between the Host and Diet setups. The shared variant appeared during EE cycle 2 (i.e., after 176 bacterial generations), reaching fixation in all Diet-evolved populations (Figure 5d, Supplementary files 7, 8), while it disappeared in the Host-evolved populations. Here, it was replaced by multiple additional variants within the same gene. At the same time, we detected mutations that exclusively occurred in each evolutionary setup. Specifically, 11 genes mutated in the Host-evolved populations at least once, while 9 exclusive mutational targets were detected in the Diet-evolved populations (Figure 5a,d, Supplementary files 7, 8). Altogether, our results show that *L. plantarum* undergoes parallel genomic evolution with and without its invertebrate animal host, further demonstrating that, contrarily to what we observed in *L. plantarum*/mouse symbiosis, in the relationship between *L. plantarum* and the fruitfly, the host nutritional environment largely dictates the microbial evolutionary trajectories both genomically and phenotypically.

## DISCUSSION

Given the complexity of host–microbial symbioses and the high variability that characterizes such associations in natural environments, understanding the evolutionary trajectories of gut microbes across different symbiotic relationships is of particular interest. In this work, we coupled experimental microbial evolution with genome sequencing and phenotypic characterization to explore the evolutionary path of *Lactiplantibacillus plantarum*, a common animal gut commensal, in vertebrate (mouse) and invertebrate (*Drosophila melanogaster*) animal models. Scientific research, including our work, demonstrated that, in the symbiosis between *Drosophila* and *L. plantarum*, the nutritional environment is the main selective agent in the evolution and adaptation of the commensal bacterium (Martino et al., 2018). Here, we asked if and how *L. plantarum* evolution varied in the context of its symbiosis with mammalian hosts, where gut commensals exhibit higher levels of colonization and mutualistic interactions are reported as more persistent than those observed in insects (Douglas, 2011; Faith et al., 2013). Although diet is well known to largely dictate gut microbes’ evolution in mammals (Carmody et al., 2015, Dapa et al, 2022, Turnbaugh et al., 2009), disentangling the respective roles of nutritional and host factors in such processes is challenging. To address this, we tracked *L. plantarum* phenotypic and genomic evolution in the intestines of mice and in their diet.

*L. plantarum* growth dynamics in the presence and absence of the mammalian animal host were similar between the two evolutionary setups (Figure 1b). We believe that, in the Diet setup, the fast and high microbial persistence can be explained by considering the structure of our experimental setting (i.e., bacterial re-inoculation at the beginning of each EE cycle), a result that was not expected in the presence of the host. Here, bacterial administration was performed at the beginning of the experimental evolution and bacteria were transferred vertically and horizontally among mice, without any further external input. In addition, the complex physical and chemical conditions encountered during transit through the mammalian gastrointestinal tract usually provide a challenge to the commensal microbiota attempting to colonize the host mucosa (Parker et al., 2018; Tannock, 2007). Here, the innate immune system of the host, the food transit, the plethora of antimicrobial compounds secreted into the gut (including hydrochloric acid, bile, hydrolytic enzymes, and antibodies), as well as other ecological factors, such as the intense microbial competition for space and nutrient resources, represent obstacles to both bacterial colonization and survival (Han et al., 2021; Louis and O’Byrne, 2010). We speculate that the high bacterial concentration observed in the Host setup is directly linked to the use of mono-colonized mice. It will be interesting to assess the *Lp* evolutionary growth dynamics in mice harboring a complex microbial community. *L. plantarum* adaptation to the mammalian host was also visible by the increased tolerance under bile acid stress and the higher persistence in the mouse intestine exhibited by the *Lp* populations evolved in the Host setup (Figure 1c,d - figure supplements 3 and 4). This suggests that, while facing the adverse conditions along the mouse GI tract, some bacterial subpopulations gradually adapted to the host intrinsic factors, resulting in improved growth, fitness, and colonization ability.

During *Lp* experimental evolution in the mouse setup, we also detected a morphological transition of bacterial colonies, characterized by the emergence of a rough phenotype that occurred both in the Host and Diet setups. Morphological transitions within a single strain population, known as pleomorphism (Wainwright, 1997), often occur in bacteria evolution and have been reported by several studies as an adaptive strategy for survival in response to fluctuating environments, especially to limiting or changing growth conditions. Moreover, *L. plantarum* had already been shown to exhibit a rough surface and cell elongation under a wide range of stress environments, including heat and ethanol shocks (Capozzi et al., 2011; Kubota et al., 2008; van Bokhorst-van de Veen et al., 2011), presence of nitrite (Wei et al., 2017), low pH (Ingham et al., 2008; Kubota et al., 2008), lactic acid stress (Pieterse et al., 2005), nutrient stress (Parlindungan et al., 2018), and bile stress (Bron et al., 2004). Interestingly, when observed with an electron microscope, the rough colonies exhibited a filamentous and chaining phenotype (Figure supplements 1b,c). In this regard, it is worth noticing that the Diet-evolved strains exhibiting the rough morphology showed a mutation in a gene encoding for the cell division protein DivIVA (Supplementary file 3). DivIVA is a coiled-coil protein first discovered in *Bacillus subtilis* and highly conserved among Gram-positive bacteria. It clusters at curved membrane areas such as the cell poles and invaginations that occur during cell division, where it serves as a scaffold protein for the recruitment of Min proteins, which spatially regulate the division process (Eswaramoorthy et al., 2011; Kaval and Halbedel, 2012). In contrast to *B. subtilis*, where the deletion of the *divIVA* gene was responsible for the formation of a filamentous and mini-cell phenotype (Cha and Stewart, 1997), ΔdivIVA mutants of *Listeria monocytogenes* exhibited a pronounced chaining phenotype (Halbedel et al., 2012). Although these cells had clearly completed cell division, they remained attached even after completion of cross-wall synthesis. This indicates that the deletion of *divIVA*, although not affecting cell division *per se*, might affect the post-divisional separation of daughter cells. However, further evidence of *divIVA* functioning within *L. plantarum* species is needed to assess our hypothesis, considering that the functionality of *divIVA* has been shown to be species-specific (Kaval and Halbedel, 2012). We next investigated the cause of the emergence of the rough morphotype. Even if we cannot assess whether the pleomorphism that emerged from both the Diet and Host-evolved rough colonies was due to the same stress factor, it is interesting to note that all rough colonies had reduced vitality and increased susceptibility to bile acids stress compared to the smooth ones, regardless of their evolutionary history (i.e., Host- or Diet-evolved) (Figures 1c, figure supplements 3,4). This is consistent with other findings, showing that the gradually increasing severity of changes in *L. plantarum* morphology coincide with a respective decrease in the bacterial growth rate (Bron et al., 2004).

Genome sequencing of *Lp* evolved with and without its mammalian host revealed weak parallelism. The Diet-evolved populations showed a significantly lower number of mutations, among which only one persisted over time (*Dipeptidase* gene). On the contrary, we detected several mammalian host-specific mutational targets, most of which are involved in carbohydrate transport and metabolism (Figure 3 and Supplementary file 4). Among these, four genes belong to the phosphotransferases system (PTS), a highly conserved bacterial phosphotransferase cascade whose components modulate many cellular functions in response to carbohydrate availability (Deutscher et al., 2006) and which has already been observed to be over-expressed or mutated in other mouse gut colonization experiments (Barreto et al., 2020; Marco et al., 2009; Zhang et al., 2013). This is in line with a recent study showing that *L. plantarum* convergently evolves across vertebrate animal hosts (i.e., human, mouse, zebrafish) by acquiring mutations primarily modulating carbohydrate utilization and acid tolerance (Huang et al., 2021).

Notably, *Lp* genomic evolution within the mouse intestine was repeatedly characterized by the emergence of hypermutators carrying multiple mutations in the *mutS, dnaE* and *dinB* genes (Figure 2b,d), which were not observed in any of the Diet-evolved populations. These genes are involved in the DNA mismatch repair system and DNA replication accuracy (Echols and Goodman, 1991; Fukui, 2010; Tompkins et al., 2003). Mutations in these regions can lead to up to a 100-fold increase in the spontaneous mutational rate compared to their wild-type counterparts (Horst, 1999; Miller, 1996; Modrich and Lahue, 1996; Sundin and Weigand, 2007) and have already been observed in clinical, environmental, and laboratory microbial populations, suggesting that the evolutionary strategies of bacteria include systems for increasing mutability (Denamur et al., 2000; Weigand and Sundin, 2012). In the context of the mammalian gut, most of the experimental research has been carried out using *E. coli*. With this species, spontaneously arising mutator bacteria can also quickly become dominant during the course of gut colonization (Barreto et al., 2020; Giraud, 2001; Giraud et al., 2008; Ramiro et al., 2020; Tenaillon et al., 2017). Such an advantage depends on the ability of the hypermutators to generate adaptive mutations rather than on the beneficial pleiotropic effects of the mutator allele, suggesting that adaptive mutations are fixed rapidly in mutator populations. Indeed, while in stable environments the maintenance of a low mutational rate is fundamental to avoid deleterious mutations that might lead to species loss, several experimental studies, performed both *in vitro* (Cox and Gibson, 1974; Mao et al., 1997) and *in vivo* (Giraud, 2001; Sniegowski et al., 1997), have shown that increased mutational rates can be beneficial to bacterial populations facing unpredictable adverse conditions, where mutations might help cells to overcome selective pressures (Bjedov et al., 2003; Denamur and Matic, 2006). At the same time, it has been shown that a high mutation rate can be initially beneficial because it allows faster adaptation, but this benefit disappears once adaptation is achieved (Giraud, 2001; Giraud et al., 2008). Although we did not assess whether the high mutation rate observed within the Host-evolved *Lp* populations is directly responsible for adaptive mutations in the mouse intestine, our results showed that the hypermutator lineages persist longer in the mouse intestine compared to the ancestral population (Figure 1d). Nevertheless, the elevated mutation rate, as well as other events commonly observed in studies of gut microbe evolution (e.g., clonal interference, epistasis, horizontal gene transfer) might mask signatures of adaptation, making it difficult to separate selection from drift (Barroso-Batista et al., 2014; Frazão et al., 2019; Giraud, 2001; Giraud et al., 2008; Zhao et al., 2019). From this standpoint, understanding the role of persisting mutations and their effect on gut microbiota fitness and physiology might be of great interest to further dissect their consequences on the evolution of the symbiosis between bacteria and its mammalian animal host.

The divergent trajectories detected by comparing *L. plantarum* evolution in a mammalian host and in its diet are in contrast with *L. plantarum* evolutionary dynamics observed in *Drosophila*, where we have previously demonstrated that gut microbes undergo parallel evolution in the presence and absence of the fruit fly (Martino et al., 2018). This indicates that selection regimes were comparable in the two environments. Specifically, in the Host setups, bacteria were horizontally and vertically transmitted among hosts (both in mice and *Drosophila*), while in the Diet setups, artificial passages of the evolving bacterial population were performed. However, we acknowledge that the experimental differences in bacterial propagation between the Host and Diet setups might not allow a direct comparison of the bacterial evolutionary dynamics.

To address this issue, in this work, we replayed *L. plantarum* experimental evolution in *Drosophila* by applying the same transferring and sampling time in both setups. This allowed us not only to demonstrate that the parallelism between *Lp* populations evolved in the presence of *Drosophila* and in its nutritional environment is repeatable, regardless of the experimental selection regimes used, but also to provide new insights into the respective role of the invertebrate host in the evolutionary path of *L. plantarum*. Indeed, we detected host-specific mutations that occurred in a later stage of bacterial evolution and showed persistence across EE cycles (Figure supplement 9). These genes mainly belong to bacterial immune evasion pathways, amino acid transport and metabolism. Specifically, *Lp* Host-evolved populations were repeatedly affected by non-synonymous mutations of the *<u>mprF</u>* gene, which encodes for an *L-O-lysylphosphatidylglycerol synthase*, an enzyme that is present in both Gram-positive and Gram-negative bacteria (Weidenmaier et al., 2003) and catalyzes the transfer of a lysyl group to the negatively charged phosphatidylglycerol (PG), a major component of the cytoplasmatic membrane (Lennarz et al., 1966; Staubitz et al., 2004). This reaction modifies the net charge of PG, neutralizing the membrane surface and thus significantly impacting the interactions with cationic antimicrobial peptides (CAMPs) produced by the host’s immune system (defensins and cathelicidins). Accordingly, the loss of Lys-PG in *mprF* mutants has been shown to lead to an increase in bacterial susceptibility to a broad variety of cationic antimicrobial peptides (Peschel et al., 2001) in different bacterial species (Samant et al., 2009; Sohlenkamp et al., 2007; Thedieck et al., 2006), thereby demonstrating a general role of mprF in bacterial immune evasion. In addition, non-synonymous Host-specific mutational targets also include *glnQ*, encoding a glutamate transport ATP-binding protein, which is involved in glutamate uptake in other Gram-positive bacteria (Capo et al., 2020), and *glnA* (encoding for a glutamine amidotransferase), involved in amino acid transport and metabolism. Specifically, glutamine amidotransferases (GATase) are enzymes that catalyze the removal of the ammonia group from a glutamine molecule and its subsequent transfer to a specific substrate, thus creating a new carbon–nitrogen group on the substrate. It is important to notice that, although such mutations were detected only in Host-evolved *Lp* populations, they ultimately did not confer a fitness advantage in the presence of the host, as Diet-evolved populations ultimately reached higher loads when associated with the fruit fly (Figure 4b). These results differ from other previous findings, according to which *Drosophila* has a positive impact on the growth of its gut microbiota (Storelli et al., 2018). To further investigate this point, we compared the growth of the two *Lp-*evolved populations in both experimental setups (Host and Diet). Notably, we found that the *Lp* concentration was always significantly higher in the presence of *Drosophila*. However, the Diet-evolved bacteria had an overall growth advantage compared to the Host-evolved populations when associated with the fruit fly (Figure 4c). Interestingly, this result was already visible from the *Lp* growth dynamics monitored during the experimental evolution (Figure 4b), where *Lp* was retrieved in a higher concentration in presence of the fly only in the early stages (∼88 *Lp* generations, EE cycle 1).

Taken together, our results suggest that the fly improves *Lp* growth in a short ecological timescale, that is in the absence of evolution, regardless of the microbial evolutionary background. On the other hand, bacterial growth is favored in the absence of its fly host in the longer term. We speculate that such different growth dynamics might be due to a combination of factors. On the one hand, the growth advantage initially conferred by the fly host to its microbiota might result in an evolutionary “relaxed” selective environment, which in turn affects bacterial growth and adaptation. On the other hand, the stronger selection occurring in the absence of the fly host might ultimately result in higher bacterial fitness in a longer timescale. Indeed, it is commonly expected that the rate of adaptation is higher when selection is stronger (Hartl and Clark, 1998). At the same time, it is worth noticing that the lower microbial load detected in the presence of the host might be partly due to the continuous ingestion of bacteria by the fruit fly, as it is known that a large fraction of ingested bacteria gets killed while passing through the stomach-like region of the *Drosophila* gut (Bing et al., 2018; Keebaugh et al., 2018; Storelli et al., 2018). However, this does not fully explain the higher *Lp* loads retrieved in the absence of the host, as it should have been visible already during the first EE cycle (Figure 4b).

Altogether, our results demonstrate that *L. plantarum* evolution diverges between insects and mammals. Specifically, we show that in *Drosophila*, the nutritional environment dictates microbial evolution, while the host benefits *L. plantarum* growth only over short ecological timescales. By contrast, in a mammalian animal model, *L. plantarum* evolution results to be divergent between the host intestine and its diet, both phenotypically (i.e., Host-evolved populations show higher adaptation to the host intestinal environment), and genomically. Here, both the emergence of hypermutators and the high persistence of mutated genes within the host’s environment strongly differed from the low variation observed in the host’s nutritional environment alone. This indicates that the mammalian animal host, together with host’s intrinsic factors, represent crucial agents of selection for the evolutionary path of gut microbes. The key questions we need to address in future studies should center on the characterization of the targets of selection, as well as the factors driving the gut microbes’ adaptation from a subspecies level to higher levels of microbial interactions. Addressing this will allow us to better understand the relationship among evolution, adaptation and microbial function and will reveal the principles that govern the colonization success, persistence, and resilience of gut microbes and how they vary across animals and humans.

## MATERIALS AND METHODS

### Bacterial strains and culture conditions

All strains used in the present study were derived from the ancestor *L. plantarum*^NIZO2877^ that was originally isolated from a sausage in Vietnam (Martino et al., 2015). At the end of each Experimental Evolution transfer or generation, the evolved strains were stored at -80°C in 1 ml of Phosphate Buffered Saline (PBS, Sigma) by adding 200 μL of 80% glycerol.

### *Drosophila* stocks and breeding

Drosophila *yw* flies were used as the reference strain in this work. Drosophila stocks were cultured at 25°C with 12/12-hour dark/light cycles on a yeast/cornmeal medium containing 50g/l inactivated yeast (rich diet) as described by Storelli et al. (Storelli et al., 2011). Poor-nutrient diet was obtained by reducing the amount of yeast extract to 8 g/l. Germ-free (GF) stocks were established and maintained as described in Storelli et al. (Storelli et al., 2011).

### *Drosophila* Diet

The fly diet used in the present study was a poor yeast diet containing 8 g inactivated dried yeast, 80 g cornmeal, 7.2 g Agar, 5.2 g methyl 4-hydroxybenzoate sodium salt, 4 mL 99% propionic acid per 1 litre. After preparation, fly food was poured in 50 mL tubes by adding 10 mL of food to each tube.

### Mouse diet

For mouse breeding and for *in vitro Lp* evolution we use mouse breeding extrudate diet V1126-000 (Ssniff, Soest, Germany). The diet was vacuum packed and sterilized by gamma-irradiation (25 kGy, Bioster, Czech Republic). It is a grain-based diet consisting of wheat, soybean products, corn (maize) products, oat middlings, minerals, soybean oil, sugar beet pu*Lp*, vitamins & trace elements, L-lysine HCl, DL-methionine. For the Experimental Transfer of *Lp*^*NIZO287*7^ in the mouse diet, the food was manually crushed, as it was initially provided in the form of pellets.

### *L. plantarum* experimental evolution in mice

Germ-free (GF) C57Bl6 mice were kept under axenic conditions in Trexler-type plastic isolators, and the absence of aerobic and anaerobic bacteria, molds, and yeast was confirmed every two weeks by standard microbiological methodology (90). The mice were kept in a room with a 12 h light-dark cycle at 22°C, fed an irradiated sterile diet V1126-000 (Ssniff, Soest, Germany) and provided sterile autoclaved water ad libitum. *L. plantarum*^NIZO2877^ was grown in De Man, Rogosa and Sharpe (Oxoid) in static culture overnight at 37°C for the monocolonization of GF mice. Seven twelve-week-old GF mice (1 male and 6 females) were colonized with a single dose (2 × 10^8^ CFU/200 μl PBS) by intragastric gavage to create F0 generation. Evolving bacteria were horizontally dispersed and vertically transmitted with no further artificial inoculation. The stability and level of colonization was checked periodically by plating of appropriate feces dilution collected from 3-4 mice on MRS agar and counting after aerobic cultivation for 48 h at 37°C (CFU/g feces). After verification of stable colonization, mice were mated and colonization of F0 and subsequent generations (F1, F2, F3, F4) was followed for 10 months. Fecal pellets were collected and pooled from 3-4 mice for the duration of the entire experimental evolution (10 months), diluted in PBS 1X and plated in MRS agar to isolate the evolving *L. plantarum* colonies in order to follow their evolution along time or stored with 20% glycerol at −80°C for future analysis. The animal experiments were approved by the Committee for the Protection and Use of Experimental Animals of the Institute of Microbiology of the Czech Academy of Sciences.

### *L. plantarum* experimental evolution in the mouse diet

The experimental evolution of *L. plantarum*^NIZO2877^ in the mouse laboratory diet was designed as follows: *Lp*^NIZO2877^ (ancestor) was cultivated at 37°C overnight in 10 mL of MRS Broth. On the following day (day 0), 1 mL of the overnight culture was centrifugated at 4000 rpm for 10 minutes and washed in sterile PBS. After proper dilutions, 10 μL of PBS-washed culture of *L. plantarum* (corresponding to 10^2^ total CFUs) were inoculated in microtubes (five total technical replicates) containing 150 mg of the crushed mouse laboratory food supplemented with 100 μL of sterile deionized water. At the same time, 100 μL of the bacterial inoculum were plated out on MRS Agar and grown at 37°C for 48 h as a control. To mimic the host’s intestine condition, bacteria were incubated at 37°C for 24 hours. On the following day (day 1), the evolved bacteria of Transfer 1 (T1) were isolated from each of the five replicates of the mouse diet. Specifically, the medium was crushed using the Tissue Lyser II (Qiagen) (frequency of 30 Hz for 40’’) in 1 mL of PBS microtubes containing 0.75/1 mm glass beads. 10 μL of the crushed medium (10^2^ total CFUs) were used to inoculate five novel sterile medium microtubes (day 0 of T2). This allowed the propagation of an evolving bacterial subpopulation derived from the ancestor on the new medium. To determine the microbial load reached at the end of the first transfer (day 1 of T1), 100 μL of the crushed medium were plated out on MRS agar at 37°C for 48 h. Each experimental Transfer followed the same experimental setup as the one described above, with the exception that, since bacteria were propagated along with the food, no further inoculation of the ancestor strain *Lp*^NIZO2877^ were performed. The EE in the mouse diet was conducted for a total of 20 Transfers.

### *L. plantarum* experimental evolution in *Drosophila*

Two EE protocols were performed simultaneously to evolve *L. plantarum*^NIZO2877^ in presence of both *Drosophila* and its diet (Host setup) or just with its diet (Diet setup). For the first generation of both setups, *L. plantarum*^NIZO2877^ was cultured overnight in 10 mL of MRS Broth at 37°C. At the same time, GF female flies had been placed inside cages containing poor nutrient GF medium to lay eggs. On the following day, 40 embryos were transferred to 5 replicate tubes containing poor nutrient diet (Host setup). 5 tubes containing poor nutrient diet were also used for the Diet setup, with the exception that no *Drosophila* eggs were added in this case. *Lp*^NIZO2877^ overnight culture was washed in sterile PBS and after proper dilutions 1 mL of PBS-washed culture was added directly on the eggs and the fly food (bacterial inoculum = 10^5^ CFU/mL). No further inoculation of the ancestor strain *Lp*^NIZO2877^ was performed after the beginning of the first generation until the end of the experimental evolution. Once the monoassociation had been performed, Host and Diet tubes were incubated at 25°C. As soon as at least 15 pupae emerged from all Host tubes, 150 mg of food were transferred from each of the ten tubes into as many new microtubes where 0.75/1mm of glass-beads were previously introduced. 1 mL of MRS broth was added in each microtube and the content was dissolved by using Tissue Lyser II (30 Hz for 1 minute). After proper dilutions, 100 μl deriving from each microtube were plated out on MRS agar plates, which were incubated at 37°C for 48h for colonies counting. Finally, 200 μl of sterile glycerol (80%) were added in each microtube to store the bacteria at -80°C. The preparation of the subsequent Host Generations (G) or Diet Transfers (T) reflected the described procedure and started from the frozen microtubes obtained in the previous generation. Depending on their bacterial concentration, an adequate number of dilutions was performed in order to inoculate 10^5^ CFU/ml to the new Diet and Host tubes for each of the following Transfers/Generations.

### Generation time of *L. plantarum*

#### Generation time of L. plantarum in Drosophila experimental setup

To determine the generation time of *L. plantarum* strains in the Drosophila Host and Diet experimental setup, we used a modified version of a method that reported the correlation between bacterial growth rate and 16S rRNA content (Poulsen et al., 1995). *L. plantarum* was cultured to stationary phase (18h) and washed in sterile PBS. Serial dilutions have been prepared and 5 μl containing a total of 10^3^ colony-forming units (CFUs) were added to 100 μl of GF poor nutrient diet with and without Drosophila larvae (Diet and Host setup respectively) and kept at 25°C. Samples were snap-frozen in liquid nitrogen at different time points across five days of growth. Bacterial RNA was extracted using NucleoSpin RNA Isolation kit (Macherey-Nagel, Germany) following manufacturer’s instructions. Reverse transcription of total extracted RNA into cDNA has been performed using Superscript II (Invitrogen, USA) according to manufacturer’s instructions. Quantitative PCR was performed in a total of 20 μl on a Biorad CFX96 apparatus (Biorad) using SYBR GreenER qPCR Supermix (Invitrogen, USA). The reaction mixture consisted of 0.5 μl of each primer (10 μM each), 12,5 μl of SYBR GreenER mix, 10 μl of water and 1,5 μl of template cDNA. The PCR conditions included 1 cycle of initial denaturation at 95°C for 2 min, followed by 45 cycles of 95°C for 10 sec and 60°C for 40 sec. Absolute quantification of 16S rRNA was conducted as follows: five 1:10 serial dilutions of the standard sample (100 ng/μl of cDNA extracted from *L. plantarum*^NIZO2877^ culture) were quantified by Real-time PCR using universal 16S primers (forward primer, UniF 5’-GTGSTGCAYGGYTGTCGTCA-3’ and reverse primer, UniR 5’-ACGTCRTCCMCACCTTCCTC-3’) (Packey et al., 2013). Each dilution has been tested in triplicate. Melting curves of the detected amplicons were analysed to ensure specific and unique amplification. Standard curves were generated plotting threshold cycle (Ct) values against the log of the standard sample amount. Based on the data obtained from the standard curve, the Ct values of the Host and Diet samples have been used to obtain the log of their 16S rRNA concentration at each time point. The 16S rRNA values during exponential phase have been used to infer the bacterial generation time following the equation reported by Widdel et al. (Widdel, 2010).

#### Generation time of L. plantarum in Mouse experimental setup

Generation time of *L. plantarum* in the mouse intestine was estimated in the jejunal loops of germ-free mice. *L. plantarum* was grown overnight in MRS broth at 37°C, centrifuged (4500rpm x 10 min), washed with sterile PBS and adjusted to 10^7^ CFU/ml. Four 8-week-old germ-free female C57Bl6 mice were anesthetized by intraperitoneal injection of ketamine/xylazine mixture. Mice were shaved on the abdomen, laparotomy was performed and 2 jejunal loops were created with nylon ligatures 10^6^ CFU of *L. plantarum* in total volume of 100 μl was applied directly into the loops using gauge needle. Intestines with loops were put back inside the abdominal cavity; mice were placed into individual boxes on heated pad (37°C) saturated with 0.5 % isofluoran. Mice were euthanized at T0 (immediate loops harvest), T1 (1 hour) and T2 (2 hours), loops were taken out from the re-opened cavity and each loop was homogenized in 1 ml sterile PBS using TissueLyser LT (Qiagen) and stainless-steel beed. Serial dilutions in PBS were plated on MRS agar and colonies were counted after 48 h at 37 °C. 10^7^ CFU/ml of *L. plantarum* in PBS and the same aliquot after incubation for 2 h at 37 °C were plated out as controls (Figure supplement 7). The experiments were approved by the Animal Care Committee of the Czech Academy of Sciences (protocol n. 18/2019) and were in accordance with the EU and NIH Guide for the Care and Use of Laboratory Animals. *L. plantarum* generation time in the mouse intestine and diet was estimated by the following formula (as reported by Kushkevych et al., 2019):

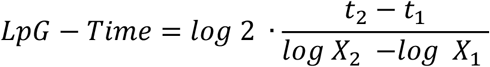

(t = time; X = colony forming units per milliliter).

### *Lp*^NIZO2877^ serial Passages in MRS broth and MRS broth + 0.3 % bile acids (BA)

The ancestral strain *Lp*^NIZO2877^ was cultured overnight at 37°C in 10 mL of MRS broth and MRS broth added with 0.3% sterilized bovine bile acids (Sigma) by using three replicates per condition. On the following day, 100 μL of the overnight culture were plated out on MRS agar plates or MRS agar plates containing 0.3% of sterile bile acids and incubated at 37°C for 48 hours for colony counting. At the same time, 10 μL of the overnight culture were transferred into a new MRS broth or MRS broth + 0.3% bile acids medium. The same procedure has been followed for 7 days, thus allowing to determine the bacterial growth (CFU/mL) overtime.

### Spectrophotometer assays

Spectrophotometer assays were carried out in presence of MRS broth or MRS broth added with 0.3% bile acids to investigate the bacterial growth by mimicking the stress conditions found in the mouse gastrointestinal tract. The strains tested are listened in Supplementary file 4. 100 μL from the - 80°C stock of each strain were cultured on MRS agar plates at 37°C for 48 hours. Next, one colony was resuspended in 60 μL of PBS and 10 μL were then tested in triplicates in a 96-wells plate containing 100 μL of MRS broth (control), or MRS broth with the addition of 0.3% sterilized bovine bile acids (Sigma), respectively. The bacterial growth was assessed turbidimetrically by measuring optical density at 600 nm every 5 minutes for 24 hours using the Multiskan™ GO Microplate Spectrophotometer (Thermo Scientific).

### Electron microscopy

Bacterial colonies transferred from cellophane into cacodylate or phosphate buffer (pH 7.2-7.4) were carefully resuspended and fixed in buffered 3% glutaraldehyde overnight at 4 °C. Thoroughly washed cells were sedimented onto poly-L-lysine coated round 12 mm coverslips for 48 h at 4 °C. Coverslips were then washed with ddH_2_O and postfixed in 1% OsO4 at room temperature for one hour. After post-fixation, the coverslips were washed three times with ddH_2_O and dehydrated in graded alcohol series (25, 50, 75, 90, 96, 100, and 100%), followed with 100% acetone, each step for 20 min. Finally, the coverslips were critical-point dried (K850, Quorum Technologies Ltd, Ringmer, UK) and sputter-coated with 3 nm of platinum (Q150T ES, Quorum Technologies Ltd, Ringmer, UK). Alternatively, pieces of cellophane with bacterial colonies were mounted onto glass slides with Scotch tape. The mounts were then put into a Petri dish with a small container filled with 2% OsO_4_ in ddH_2_O. Fixation in osmium vapor was then performed in closed Petri dishes in the desiccator for several days at room temperature. Pieces of cellophane with fixed colonies were mounted onto standard aluminium stubs and sputter-coated with 3 nm of platinum (Q150T ES, Quorum Technologies Ltd, Ringmer, UK). All samples were examined in an FEI Nova NanoSEM scanning electron microscope (FEI, Brno, Czech Republic) at 3 to 5 kV using ETD, CBS, and TLD detectors.

### Adaptation of *L. plantarum* to the mouse intestine

Three C57Bl6 mice (2 month-old) were colonized with a single dose (2 × 10^8^ CFU/200 μl PBS) of the *Lp*^NIZO2877^ ancestral strain and *Lp*^NIZO2877^-derived population evolved in the mouse intestine (sample: F0-10 months) by intragastric gavage. Fecal pellets were sampled every 12 hours (until 72 hours after gavage) and immediately freezed at -80°C. Bacterial DNA was extracted from mice stool using NucleoSpin DNA Isolation kit (740472.50 Macherey-Nagel, Germany) following manufacturer’s instructions. Real-time PCR amplifications were performed on a LightCycler 480 thermal cycler (Roche Diagnostic, Mannheim, Germany) in a final volume of 10 µl, which included 2.5 µl of DNA. The PowerUp™ SYBR™ Green Master Mix (Applied Biosystems™, USA) was used together with 0.25 µl of each primer. Primers designed on *L. plantarum ackA* gene were used to specifically amplify *L. plantarum* DNA, while universal 16S primers were used to amplify the total bacterial DNA (Supplementary File 7). The cycling conditions were as follows: 50 °C for 2 min, followed by 2 min at 95 °C, and 45 cycles at 95 °C for 10 s and 60°C for 1 min. Outputs of real-time amplifications were analysed by means of the LightCycler 480 Basic Software Version 1.2 (Roche Diagnostic, Mannheim, Germany). The amount of *L. plantarum* DNA detected was normalized to the total bacterial DNA values to account for DNA extraction efficiency according to cycle threshold analysis (ΔC_T_).

### Fitness assessment of *L. plantarum*-evolved populations in *Drosophila* Host- and Diet-setups

*Lp*-Host and Diet-evolved populations belonging to *Drosophila* Generation/Transfer17 were tested. Ten independent replicate populations (Host: five replicates, Diet: five replicates) were analysed. Specifically, 10^4^ CFU/mL of bacteria taken from each *Lp*-evolved population were inoculated into new microtubes (N = 10 microtubes per evolutionary background) containing 250 μL of *Drosophila* food. One-day old *Drosophila* larvae were added to five out of the 10 microtubes (Host setup). The remaining five microtubes, which included only the fly diet, represented the Diet setup. After 11 days of growth at 25°C, the whole content was transferred from each sample into novel microtubes where 0.75/1mm of glass-beads and 1 mL of MRS broth were previously introduced and the content was dissolved by using Tissue Lyser II (30 Hz for 1 minute). After proper dilutions, 100 μL from each microtube were plated out on MRS agar plates and cultured at 37°C for 48 h for colony counting.

### DNA extraction and PCR amplification of *mutS* gene

Bacterial DNA was extracted from mice stool using NucleoSpin DNA Isolation kit (740472.50 Macherey-Nagel, Germany) following manufacturer’s instructions. Primers for the *mutS* gene were designed from the most conserved regions by using Primer3 software (http://frodo.wi.mit.edu/primer3/), with a length of 19 to 25 nucleotides (Supplementary file 7). The PCR amplifications were performed in a Euroclone One Advanced thermal cycler (Celbio, Milan, Italy). The PCRs were performed in a final volume of 20 μl of amplification mix containing 1 U of GoTaq polymerase (Promega, Madison, WI), 1× GoTaq buffer, 1.5 mM MgCl2, 0.2 mM each deoxynucleoside triphosphate (dNTP), 250 mM each primer, and 5 ng of genomic DNA as the template. Amplified products were analyzed by electrophoresis on 1.8% agarose-Tris-acetate-EDTA (TAE) gels, stained with SYBR Safe (Invitrogen, Carlsbad, CA), and visualized on a UV transilluminator. Bidirectional sequencing of target gene was performed using the respective primer pairs used for PCR amplifications as sense and antisense sequencing primers by BMR Genomics, Padua, Italy.

### DNA extraction and whole-genome sequencing

The bacterial samples processed for whole genome sequencing are listened in Supplementary file 1. For the bacterial populations evolved in the mouse intestine, fecal pellets pooled from 3-4 mice were analysed per time point. For the DNA extraction, 100 μL of each bacterial population or strain have been plated out in two MRS agar plates and incubated for 48 h at 37°C. Genomic DNA was extracted from a mixture of > 1000 colonies per sample by using the Power Soil DNA extraction Kit (Qiagen) by following manufacturer’s instructions. DNA library construction and sequencing were carried out by the EMBL Genomics Core Facilities (Heidelberg, Germany). Each sample was pair-end sequenced on an Illumina MiSeq Benchtop Sequencer. Standard procedures produced data sets of Illumina paired-end 250 bp read pairs. The Raw data (FASTQ) have been deposited at NCBI under the temporary Accession number SUB11111164.

### Mutation identification

Raw reads were trimmed and filtered using the parameter SLIDINGWINDOW in Trimmomatic (Bolger et al., 2014), with a 4-base wide sliding window and cutting when the average quality per base drops below 20. The mean coverage per population was between 124 and 193. Processed reads were aligned and analyzed against their respective reference strain (ancestor) genome (*Lp*^NIZO2877^) (accession number LKHZ01000000). Candidate mutations were identified by running two passes of the *breseq* pipeline in polymorphism mode (Deatherage and Barrick, 2014). Initially, each sample was analyzed individually. Mutation predictions that passed default filtering cutoffs in any one sample were merged into one overall list of candidates. Then, *breseq* was rerun a second time on each sample with the combined list as a user input file so that it output the counts of reads supporting the mutant and reference alleles for each mutation in all samples, even when a potential mutation was at a low frequency or did not pass other default filtering thresholds in a given sample. We further filtered this list of candidate mutations to remove false-positive calls, including those caused by reads that were mapped incorrectly due to incomplete assembly of the reference genome, and to distinguish low-frequency mutations from sequencing errors. For each candidate mutation, we first used examined a Poisson model of the counts of reads supporting the mutant allele relative to the total counts of reads supporting either the mutant allele or the reference allele in each sample. Real mutations that are sweeping through populations should have frequencies that significantly deviate in some samples from the background distribution across all samples. To test for this signal, we calculated p values for the significance of sample as a fixed factor in the Poisson model for each mutation. Candidate mutations with p values that were not significant after correcting for multiple testing using the Benjamini-Hochberg procedure with a false-discovery rate of 5% were removed. We next eliminated candidate mutations with frequencies that were ≥50% in at least half the samples from all treatments. Then, we kept only mutations that had a >20% range in their predicted frequencies among the samples in a given evolution treatment or that both reached a frequency of >10% and appeared in less than or equal to half of these samples to arrive at the final lists of mutations that were analyzed.

### Mutation accumulation rates and spectra

We fit linear models to the summed frequencies of all mutations observed in each sample to estimate average rates of mutation accumulation per generation. These models were constrained to have no initial mutations (i.e., an intercept term of zero). For the fly diet, fly host, and mouse diet treatments we fit models with one rate of mutation accumulation. For the mouse host treatment, we fit three rates: one for ancestral (nonmutator) lineages, one for *mutS* A41T lineages, and one for *mutS* Δ1303 lineages. We modeled the total mutation frequency in a sample as the sum of each of these three population’s rates multiplied by its frequency in the population and the number of generations that had elapsed from the beginning of the experiment up to that sample. This procedure assumes that both mutator lineages evolved close to the beginning of the experiment and experienced one constant rate of mutation accumulation throughout their history. The frequencies of the two *mutS* alleles in each sample were used to estimate the fraction of the population in each of the three categories. The *mutS* lineage frequencies were normalized to a total of 100% when they slightly exceeded this value when added together due to experimental and/or sampling errors in the estimates of allele frequencies from sequencing reads.

### Muller plots

We included only mutations that reached at least a 10% frequency in a population when constructing Muller plots and assumed there was no recombination between bacteria during the experiment (i.e., purely asexual reproduction). Since whole-population (metagenomic) sequencing does not provide information about linkage between mutations, we inferred which mutations were likely in the same genetic backgrounds from how their frequencies changed over time. We also assumed that once a lineage had a mutation in a certain gene that it was unlikely to sustain a second mutation in the same gene since these populations retained the low ancestral mutation rate. These rules disambiguated how most mutations could be ordered. We then corrected the genotype frequencies observed at each time point for two types of errors resulting from how the frequencies of all mutations in a sample are estimated independently from the sequencing reads overlapping each genomic site. First, we reduced the frequencies of genotypes with new mutations that exceeded the frequencies of their ancestral genotypes to fit within the earlier group. Second, we normalized the total frequencies of all genotypes at a given time point to 100% if it exceeded this value. Both types of corrections changed the inferred genotype frequencies by <10% in all cases. We used the ggmuller R package (Noble, 2019) to plot the resulting dynamics and then manually adjusted the locations of the curves between the time points with measured values to improve the visibility of mutations and lineages.

### Data analysis

Data representation and statistical analyses were performed using GraphPad PRISM 9 software (GraphPad software, www.graphpad.com). All the pairwise comparison were performed by using the unpaired t-test (*p≤0.05, **p<0.01, ***p<0.001). *Drosophila* developmental time, mid-point of *Lp* exponential growth phase (bile acids test) and ANCOVA test between *Lp* Host and Diet-growth trend of *Drosophila* setup have been analyzed through dedicated scripts on R Studio software (RStudio Team, www.rstudio.com) (significance p <0.05; *p<0.05, **p<0.01, ***p<0.001). Predicted gene functional categories have been determined according to their COG group with EggNOG v.5.1 (Huerta-Cepas et al., 2016).

## ACKNOWLEDGEMENTS

We would like to thank Jaroslava Valerová and Šárka Maisnerová for excellent technical assistance. Research in MEM lab was supported by the STARS@UNIPD Funding Programme (Starting Grant 2017) of the University of Padova and by the Visiting Programme 2018 grant from Fondazione Cassa di Risparmio di Padua e Rovigo (CARIPARO Foundation). MS lab was supported by the Czech Science Foundation JUNIOR STAR grant (GAČR 21-19640M) and Ministry of Education, Youth and Sports of the Czech Republic (EMBO Installation Grant 4139). JEB lab was supported by the Welch Foundation (F-1979-20190330), the U.S. National Science Foundation (DEB-1813069), and the U.S. Army Research Office (W911NF-20-1-0195). The authors acknowledge the Texas Advanced Computing Center (TACC) at The University of Texas at Austin for providing high-performance computing resources.

## COMPETING INTERESTS

The authors declared no competing interest.

## FOOTNOTES

Author contributions: M.E.M. and M.S. designed research; E.M., M.G, D.S., A.J., O.B., O.K., N.S., T.H., M.S. performed research; E.M., I.G., J.B., M.S., M.E.M. analyzed data; and E.M., M.S., M.E.M. wrote the paper.

---

**Supplementary Figure 1.**
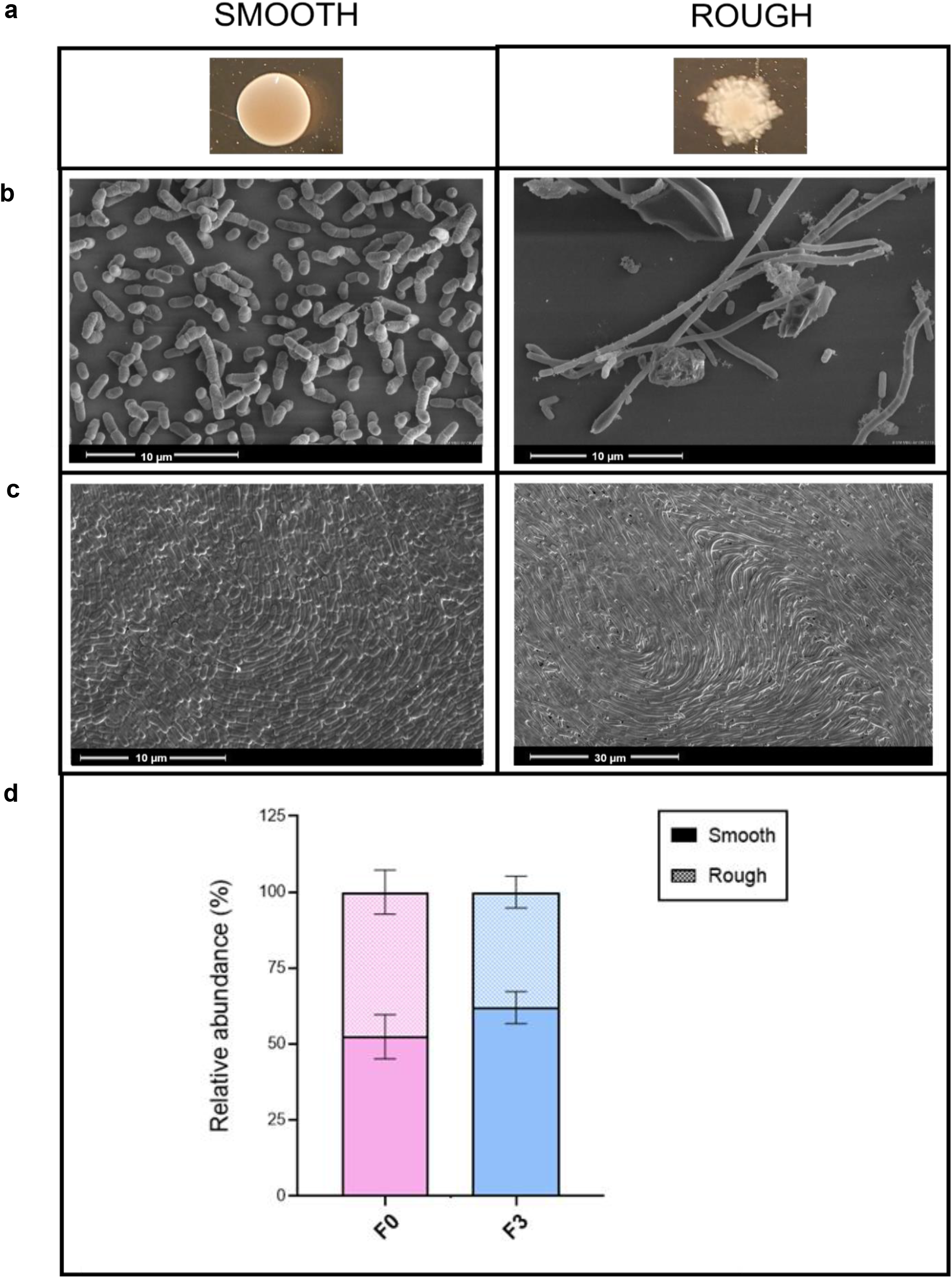
A) Macroscopical appearance of smooth/rough morphotypes isolated during Lp evolution in the mouse intestine. (B and C) Microscopical appearance of smooth/rough Lp morphotypes at the electron microscope. (C) Relative abundance of smooth and rough Lp morphotypes observed at mice generations 0 (F0) and 3 (F3). Lines above each bar indicate the standard error of the mean (SEM) determined by considering three replicates for each generation.

**Supplementary Figure 2.**
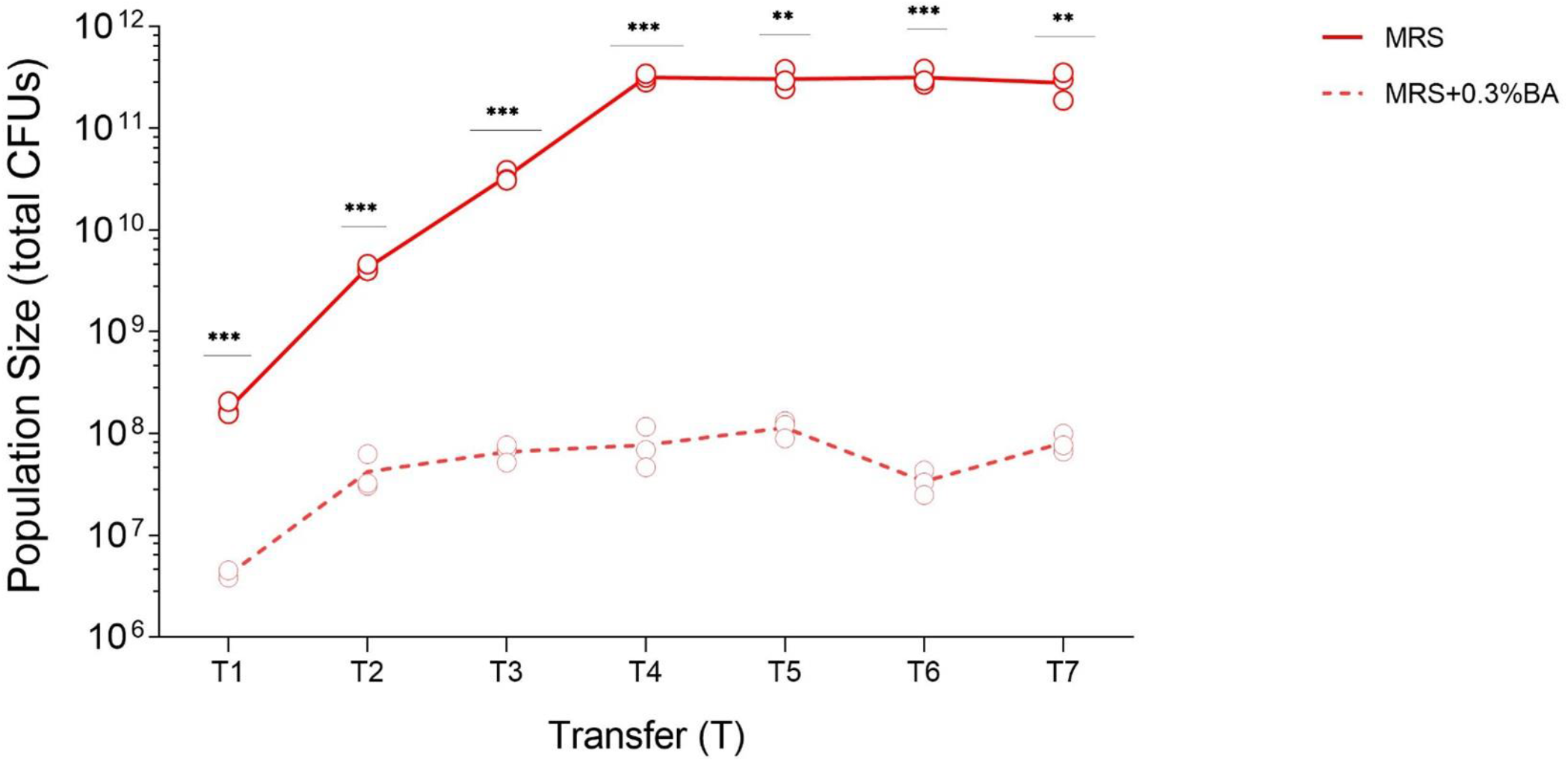
*Lp*^NIZO2877^growth monitored during serial Transfers (T) in MRS broth and MRS broth added to with 0.3% bile acid (BA). At each transfer, the three circles represent the growth obtained from each of the three experimental replicates. Asterisks refer to statistical comparison between bacterial CFUs obtained from the two experimental conditions at each transfer (unpaired t-test; * p ≤ 0.05, ** p < 0.01, and *** p < 0.001).

**Supplementary Figure 3.**
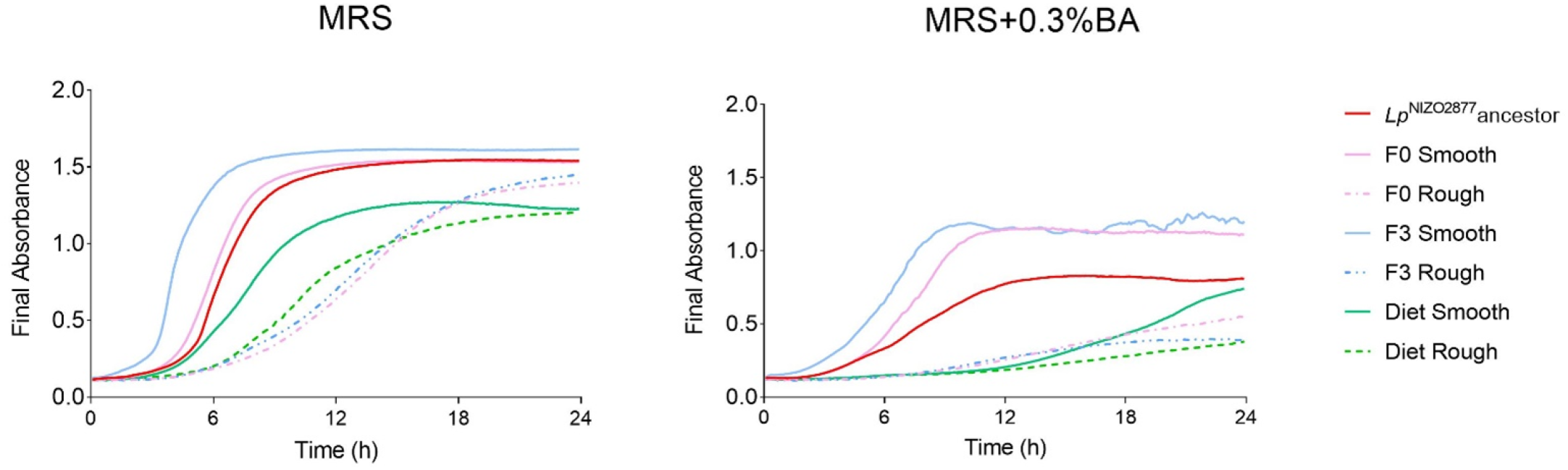
Growth curves of the *Lp* strains under standard growth conditions (MRS broth) and in MRS broth added to with 0.3% bile acid (BA). Each curve represents the mean of at least three replicates.

**Supplementary Figure 4.**
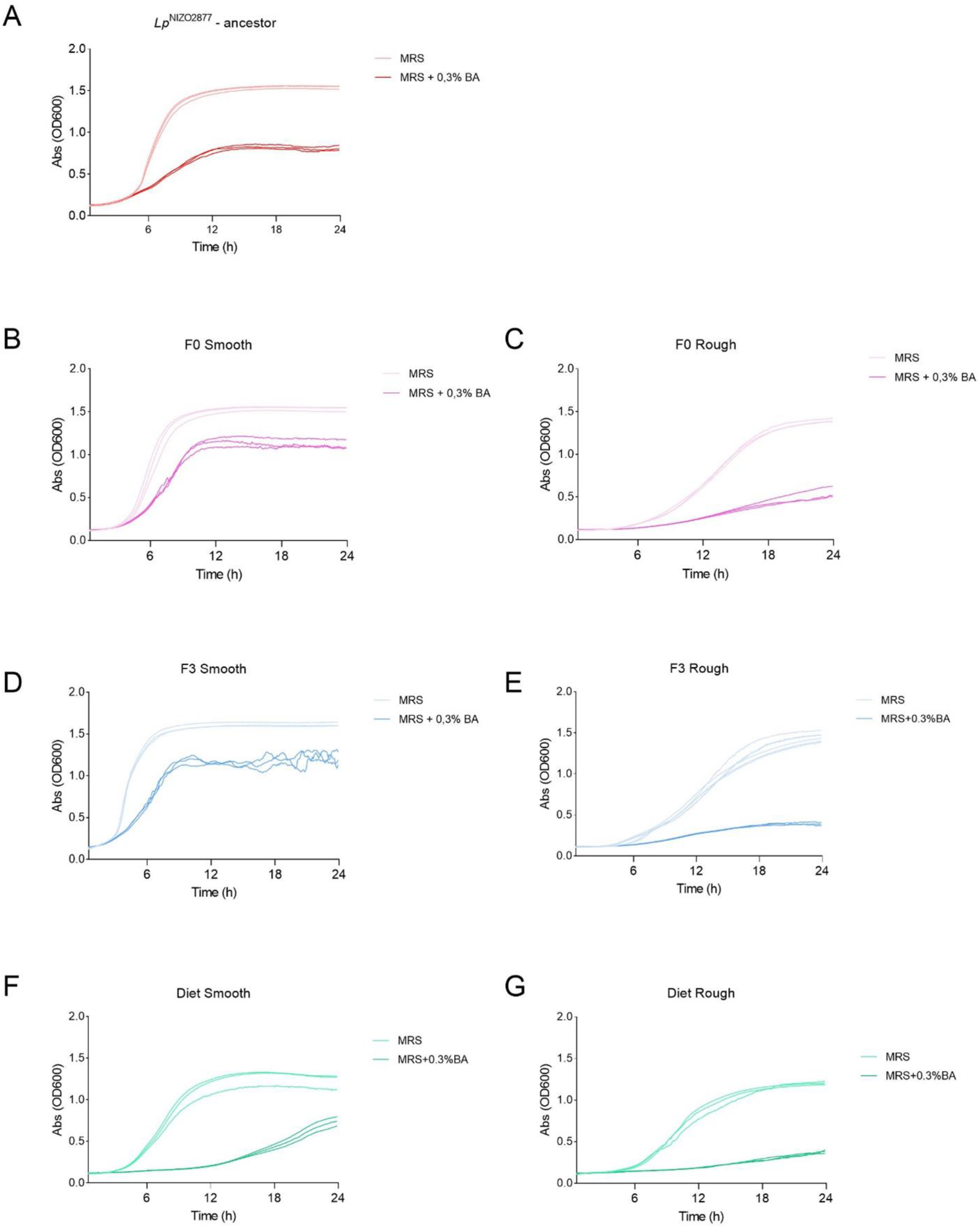
Standard growth curves of the *Lp* strains cultured in MRS broth and MRS broth + 0.3% bile acid (BA). The strains tested include (A) the *Lp*^NIZO2877^ancestor; (B, C) Smooth and Rough colonies isolated from mice generation 0; (D, E) Smooth and rough colonies isolated from mice generation 3; (F, G) Smooth and Rough colonies isolated from the Diet setup.

**Supplementary Figure 5.**
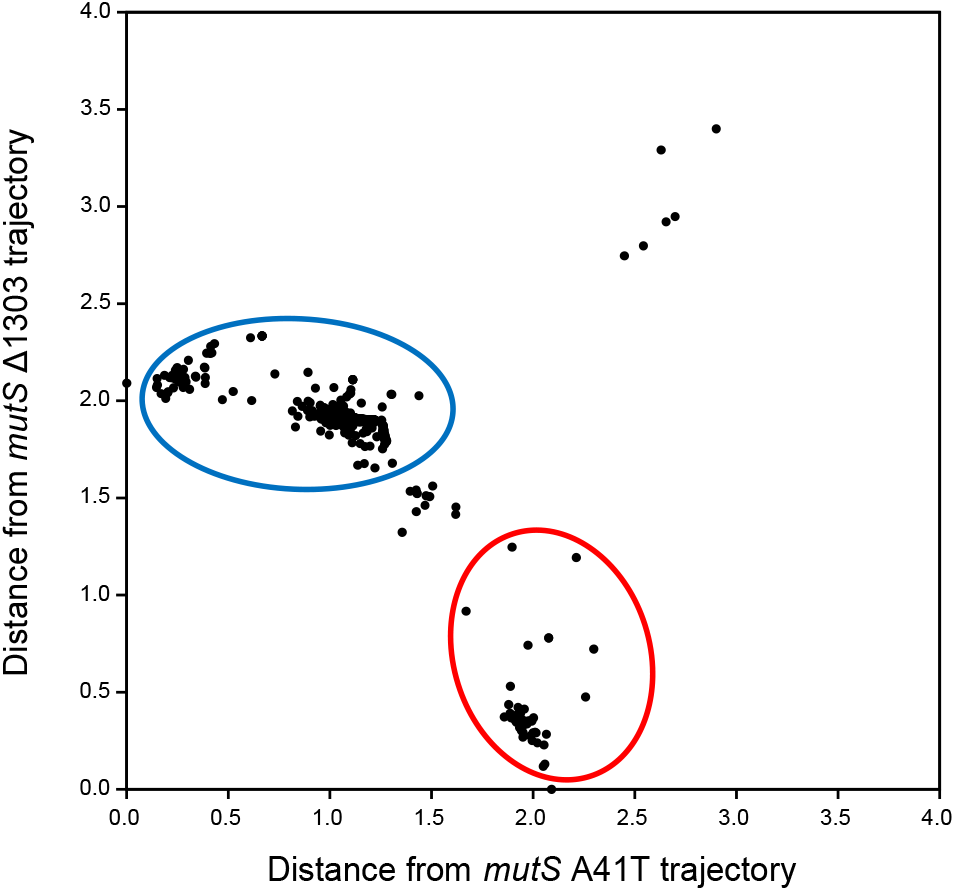
Total Manhattan distances between the frequencies of different mutations and the *mutS* A41T and *mutS Δ*1303 alleles over all samples from the mouse evolution experiment were used to classify mutations as occurring in each hypermutator lineage for examining the base substitution spectra. Mutations with distances < 0.4 to both lineages rose to a high frequency along with each *mutS* mutation. The cluster of mutations at a distance of ∼1.0 from *mutS* A41T swept within this lineage later in the experiment.

**Supplementary Figure 6.**
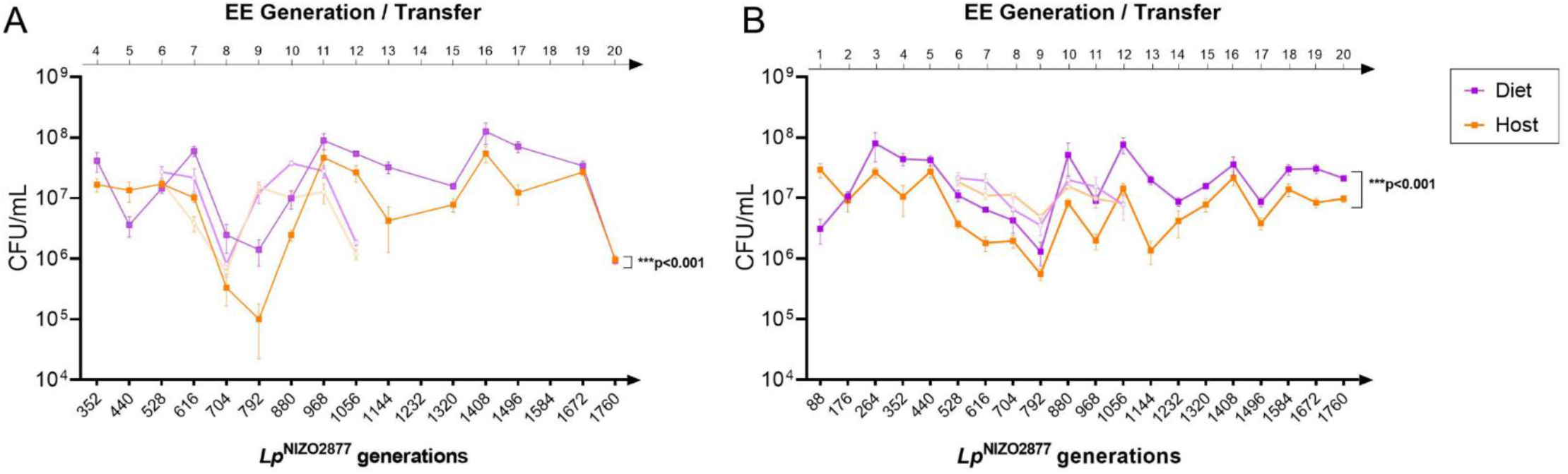
Bold full lines indicate *Lp*^NIZO2877^growth monitored after 7 (A) and 11 days (B) of incubation in the presence (Host setup) or absence (Diet setup) of Drosophila. Lighter full lines indicate the re-monitoring of *Lp* growth from Generations/Transfers 6 to 12. ANCOVA (* p < 0.01, ** p < 0.001, and *** p < 0.0001).

**Supplementary Figure 7.**
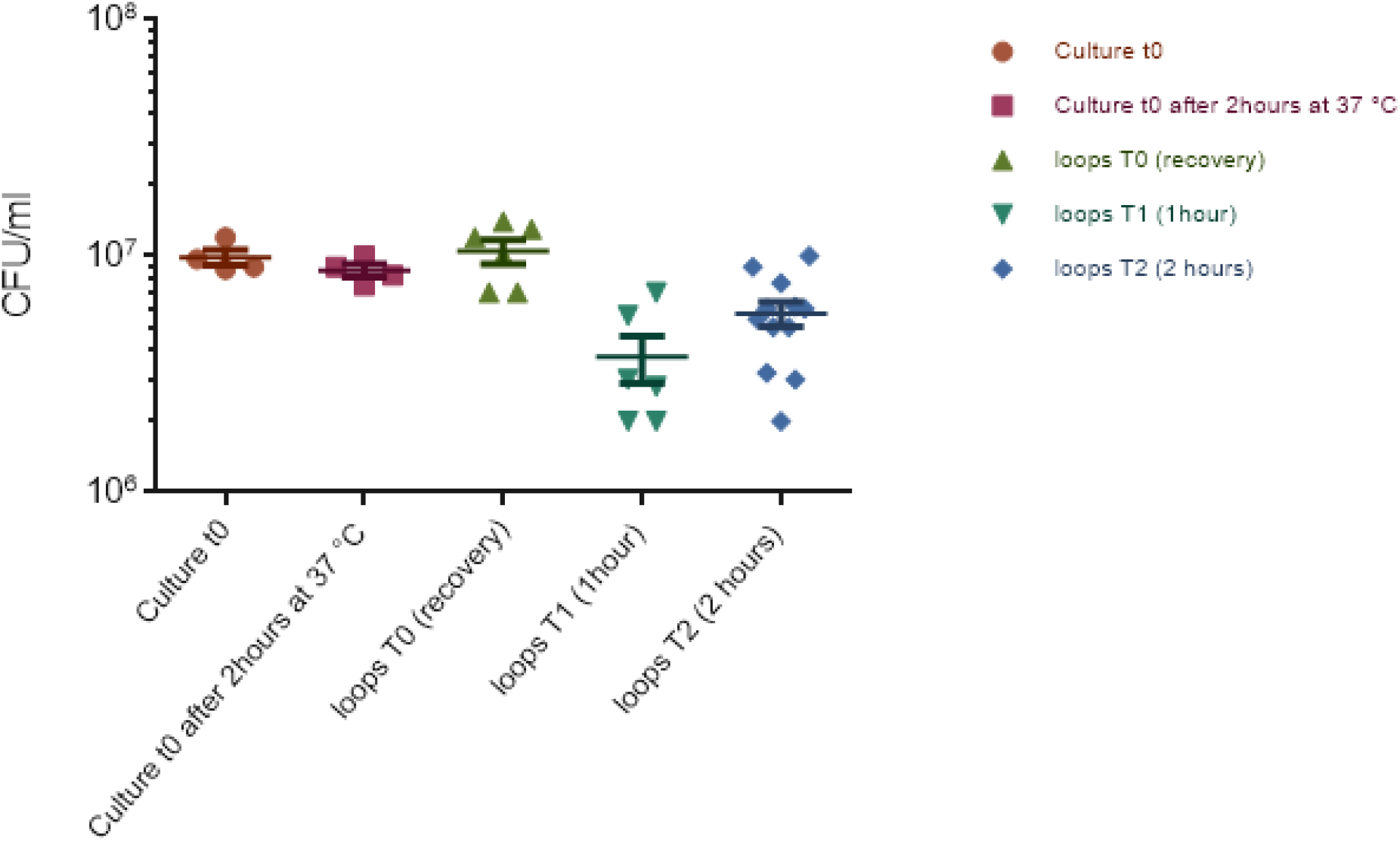
*L. plantarum* loads retrieved in the jejunal loops of germ-free mice. Bars indicate the standard error of the mean (SEM).

## Notes

### Competing Interest Statement

The authors have declared no competing interest.

